# Isoform-resolved genome annotation enables mapping of tissue-specific betalain regulation in amaranth

**DOI:** 10.1101/2023.08.24.554588

**Authors:** Tom S. Winkler, Susanne K. Vollmer, Nadine Dyballa-Rukes, Sabine Metzger, Markus G Stetter

## Abstract

• Betalains are coloring pigments produced in some families of the order Caryophyllales, where they replace anthocyanins as coloring pigments. While the betalain pathway itself is well studied, the tissue-specific regulation of the pathway remains mostly unknown.

• We enhance the high-quality *Amaranthus hypochondriacus* reference genome and produce a substantially more complete genome annotation, incorporating isoform details. We annotate betalain and anthocyanin pathway genes along with their regulators in amaranth and map the genetic control and tissue-specific regulation of the betalain pathway.

• Our improved genome annotation allowed us to identify causal mutations that lead to a knock-out of red betacyanins in natural accessions of amaranth. We reveal the tissue-specific regulation of flower color via a previously uncharacterized MYB transcription factor, *AhMYB2*. Downregulation of *AhMYB2* in the flower leads to reduced expression of key betalain enzyme genes and loss of red flower color.

• Our improved amaranth reference genome represents the most complete genome of amaranth to date and a valuable resource for betalain and amaranth research. High similarity of the flower betalain regulator *AhMYB2* to anthocyanin regulators and a partially conserved interaction motif support the co-option of anthocyanin regulators for the betalain pathway as possible reason for mutual exclusiveness of the two pigments.

## Introduction

Coloring pigments fulfill various functions in plants, including UV protection and pollinator attraction (Koes *et al*. 1994; Młodzińska *et al*. 2009). While anthocyanins are commonly observed as red pigments, the functionally similar but biochemically distinct betalains function as pigments in many families of the Caryophyllales (Brockington *et al*. 2011; Jain and Gould 2015; Lopez-Nieves *et al*. 2018; Sunnadeniya *et al*. 2016). Like anthocyanins, betalains have been associated with photoprotection, abiotic stress resistance and antioxidant activity (Jain and Gould 2015). It is noteworthy that no plant species producing both anthocyanins and betalains has been identified, suggesting a replacement of widespread anthocyanins by betalains in multiple lineages of the Caryphyllales (Brockington *et al*. 2015; Timoneda *et al*. 2019). However, the transition and evolution of the mutually exclusive presence of these two pigment pathways remains poorly understood.

Reference genome assemblies enable the identification of functional elements in genomes and allow to understand the evolution of these elements (Kersey 2019). In particular, high-quality gene annotations can give insights into gene families, biosynthetic pathways and provide an invaluable resource to study gene expression patterns and genotype-phenotype associations. Still, the annotation and quantification of isoforms is challenging for computational annotation (Salzberg 2019). The inclusion of full-length transcript sequencing data improves accuracy and provides information about splicing-variation (Xia *et al*. 2019; Wang *et al*. 2016). Sequencing and assembling genomes are becoming common tasks and well-annotated reference genomes are available for most model plants. However, these resources are still scarce for non-model species. As most betalain producing plants are non-model species, only few well-annotated reference genome sequences are available (Dohm *et al*. 2014; Wang *et al*. 2023; Chen *et al*. 2021), hindering the study of the regulation of those pigments.

The anthocyanin biosynthesis pathway, as part of the flavonoid pathway, has been studied in detail and is well understood despite its high complexity and tissue-specific regulation (Fig. 1a; Deng and Lu 2017; Xu *et al*. 2015; Lai *et al*. 2013). Betalains and particularly their transcriptional regulation on the other hand, have received far less attention. Both tyrosine, as basis of the betalain pathway, and phenylalanine, as initial substrate of the anthocyanin pathway, are derived from the same initial substrate in arogenate, but their downstream pathways are fundamentally different (Młodzińska *et al*. 2009; Lopez-Nieves *et al*. 2018). Betalain production only requires the activity of CYP76AD and DODA genes to produce yellow betaxanthins (Fig. 1b; Hatlestad *et al*. 2012; Polturak *et al*. 2016, 2018; Sunnadeniya *et al*. 2016; Christinet *et al*. 2004). The activity of specialised CYP76AD enzymes and the subsequent glucosylation through glucosyltransferases allow the synthesis of red betacyanins (Polturak *et al*. 2016; Sunnadeniya *et al*. 2016; Brockington *et al*. 2015; Sasaki *et al*. 2004; Vogt *et al*. 1999; Vogt 2002). Despite the characterization of flavonoid and betalain pathway genes in different species (Chang *et al*. 2021; Zheng *et al*. 2016; Casique-Arroyo *et al*. 2014), understanding of the conservation and regulation of both pathways remains limited.

**Figure 1.**
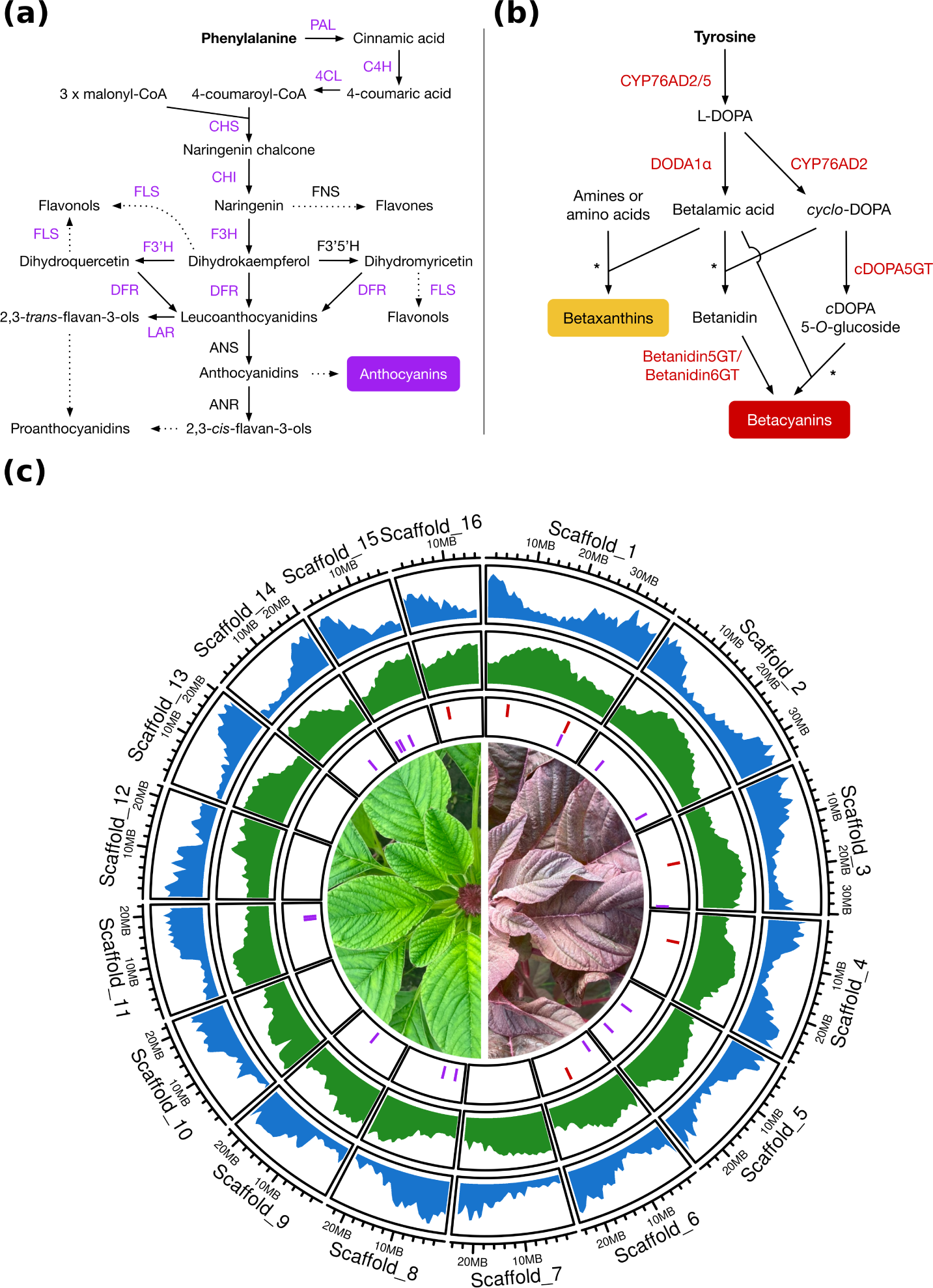
Annotation of the *A. hypochondriacus* genome v2.2. (a) Simplified schematics of the general phenyl-propanoid and flavonoid, and (b) betalain pathways. Colored pathway genes were identified in the *A. hypochondriacus* genome. (c) Circos plot representation of genome annotation v2.2 depicts gene density (blue) and repetitive element density (green) calculated in 1 Mb windows along the genome. The genomic positions of general phenylpropanoid and flavonoid (purple) and betalain pathway genes (red) were annotated in the inner track (colors as in a and b). Sample pictures of green and red colored amaranth accessions into the centre.

Although the main enzymes of the betalain pathway have been identified in recent years (Hatlestad *et al*. 2012; Christinet *et al*. 2004; Sasaki *et al*. 2004; Vogt *et al*. 1999; Vogt 2002), only few betalain regulators, including MYB transcription factors, have been described (Hatlestad *et al*. 2015; Xie *et al*. 2021, 2023; Zhang *et al*. 2021; Zeng *et al*. 2023). MYB transcription factors are best known for their involvement in the regulation of the flavonoid and anthocyanin biosynthesis pathway (Deng and Lu 2017; Li 2014; Gonzalez *et al*. 2008; Lloyd *et al*. 2017). The MYB transcription factors can be classified into subfamilies based on the number of adjacent repeats of the MYB DNA-binding domain (Jin and Martin 1999). Further classification of the R2R3 subfamily into conserved subgroups has been used to transfer experimentally validated regulatory functions to different species (Stracke *et al*. 2001; Yanhui *et al*. 2006; Stracke *et al*. 2014). Conserved subgroups participate in the regulation of the flavonoid biosynthesis in *Arabidopsis thaliana* (Stracke *et al*. 2007), grape (Kobayashi *et al*. 2002), strawberry (Schaart *et al*. 2013) and many other plant species (Naik *et al*. 2021; Lai *et al*. 2013; Brendolise *et al*. 2017). The MYB transcription factor that has been shown to regulate betalain biosynthesis in *Beta vulgaris* leaves was most similar to anthocyanin regulating transcription factors from other species (Hatlestad *et al*. 2015). This finding suggested the co-option of anthocyanin regulating transcription factors for the regulation of the chemically distinct but functionally similar betalain pathway (Hatlestad *et al*. 2015; Lloyd *et al*. 2017). However, the genetic basis and evolutionary history of this co-option remains unclear.

In this study, we generate a new reference genome for the betalain producing pseudo-cereal crop, amaranth (*A. hypochondriacus* L.). The crop forms a species complex with two other grain crops (*A. caudatus* and *A. cruentus*) and two wild relatives (*A. hybridus* and *A. quitensis*), which share a complex domestication history (Stetter and Schmid 2017; Gonçalves-Dias *et al*. 2023). We remove sequencing errors from the previous assembly (Lightfoot *et al*. 2017) and combine *ab initio* prediction and long-read isoform sequencing data to create a new, highly complete genome annotation for grain amaranth. We use this novel genome annotation to identify and annotate betalain and flavonoid color pathway genes in the genome of *A. hypochondriacus*. To identify putative regulators of these pigment pathways, we identify MYB transcription factors genome-wide and functionally assign them into conserved subgroups. Using our phylogenetic analysis, we identify multiple candidate MYBs for betalain pathway regulation in amaranth. We utilize different crosses to map genomic regions regulating amaranth tissue color and identify causal variants for color loss in a betalain pathway gene and the flower color regulator *AhMYB2* on color QTLs. *AhMYB2* has high similarity to anthocyanin regulators and regulates multiple enzymes of the betalain pathway. Understanding the tissue-specific regulation of betalain biosynthesis in amaranth can help to elucidate the evolution of this novel pigment pathway and the mutually exclusive nature of betalains and anthocyanins.

## Materials and Methods

### Improving the A. hypochondriacus reference genome

We first corrected indel errors in the *A. hypochondriacus* L. reference assembly version 2.1 (Lightfoot *et al*. 2017), using NextPolish (Hu *et al*. 2020, Supplementary Methods S1). To polish the assembly, we used the publicly available whole-genome paired-end sequencing reads from the second *A. hypochondriacus* genome assembly (Bioproject ID: PRJNA290113, SRA ID: SRR2106212) (Lightfoot *et al*. 2017). To prepare the whole-genome sequencing (WGS) reads for genome polishing with NextPolish (Hu *et al*. 2020), we removed all read pairs containing ambiguous bases using *bbduk.sh* from the BBTools suite v38.94 (https://jgi.doe.gov/data-and-tools/ software-tools/bbtools/). We annotated repetitive elements in the polished genome using Repeatmodeler v2.0.1 (Flynn *et al*. 2020, Supplementary Methods S1). We identified organellar contamination within the assembly by mapping the *B. vulgaris* mitochondrial genome (Kubo *et al*. 2000) and the *A. hypochondriacus* chloroplast genome (Chaney *et al*. 2016) against the polished assembly using minimap2 v2.17 (Li 2018).

### Plant material and genome annotation datasets

To guide computational prediction and augment the *ab initio* gene prediction, we used short- and long-read RNA sequencing data. Both short- and long-read RNA sequencing data stems from the same accession (PI 558499) that was used to generate the *A. hypochondriacus* reference genome v2.1 (Lightfoot *et al*. 2017). The short-read RNA-seq data from eight different tissues and conditions from PI 558499 was obtained from the publicly available dataset (BioProject Accession PRJNA263128) generated by Clouse *et al*. (2016). The RNA-seq dataset consisted of 352,557,994 reads spanning a total of 31.7 Gb (average of 3.97 Gb per tissue type, 90 base pair (bp) paired-end HiSeq Illumina sequencing reads; Clouse *et al*. 2016).

For the generation of long-read RNA sequencing data, material was harvested from root, cotyledons, flower, pollen, and developing and mature seeds of *A. hypochondriacus* accession PI 558499. Root tissue and cotyledons of approximately 20 seedlings were collected 3 days after germination on wet filter paper. All other tissues were collected from one single plant, grown under controlled short-day conditions. Night and day temperatures were set to 19 *°C* and 24 *°C*, respectively. Plants were grown at 60 % air humidity under well-watered conditions. Leaf tissue was collected from a fully developed young leaf at the beginning of flowering. Flower tissue was collected from unopened flowers at the tip of the inflorescence. Pollen was collected by shaking the inflorescence inside a paper bag during full bloom. Pollen collection was repeated three times within one week. Developing seeds were extracted from the flower with forceps to avoid sampling flower tissue. Approximately 20 young seeds were collected at full size, but still soft. Mature seeds were fully ripe and stored before RNA extraction. All other samples were flash frozen in liquid nitrogen and stored at -80 *°C* until RNA extraction. We extracted RNA using the Invitrogen PureLink Plant RNA Reagent (Thermo Scientific™, Thermo Fisher Scientific Inc., Massachusetts, USA) following the manufacturers protocol. PacBio isoform sequencing (Iso-Seq) was conducted by the West German Genome Center (WGGC) using the extracted RNA samples from different tissues (Table S1).

### Genome annotation

Iso-Seq circular consensus sequencing (CCS) reads were processed using Isoseq3 v3.3.0 (https://github.com/ PacificBiosciences/IsoSeq; Table S2) and cDNAcupcake v28.0.0 (https://github.com/Magdoll/cDNA_Cupcake) and corrected with the polished reference genome sequence using SQANTI3 v4.2 (Tardaguila *et al*. 2018, Supplementary Methods S2). To assess the effect of genome polishing on the assembly of full-length Iso-Seq transcripts, we also processed the reads using the unpolished reference genome (Lightfoot *et al*. 2017). We predicted open reading frames (ORFs) in the transcripts using CPC2 (Kang *et al*. 2017) and evaluated completeness of the transcript set using BUSCO (Simão *et al*. 2015) in protein mode. We used BRAKER2 v2.1.6 (Brůna *et al*. 2021) for *ab initio* gene prediction using the softmasked *A. hypochondriacus* genome assembly v2.2 (Supplementary Methods S3). Since it was unlikely for real genes in the genome to be covered neither by RNA sequencing reads nor by protein alignments and these genes contributed only marginally to the BUSCO score, we excluded predicted genes without external support from the annotation. In order to combine the computational prediction of gene structures of BRAKER2 with sequenced full-length transcripts, we used TSEBRA v1.0.3 (Gabriel *et al*. 2021) to select gene structure annotations from full-length transcripts and the BRAKER2 annotation based on their external support (Supplementary Methods S4). We assessed completeness of the genome annotation v2.2 using the annotated protein sequences as input for BUSCO v5.2.2 (Simão *et al*. 2015).

Functional annotation of annotated protein sequences was carried out using eggNOG-mapper v2.17 (Cantalapiedra *et al*. 2021), Mercator4 v.5.0 (Schwacke *et al*. 2019) and Interproscan v5.56-89.0 (Jones *et al*. 2014), retrieving pathway and gene onthology (GO) annotations (Supplementary Methods S5). We identified betalain pathway genes using BLAST (Altschul *et al*. 1997) searches with previously described betalain pathway genes (Table S3, Supplementary Methods S5). We used the flavonoid pathway identification pipeline KIPEs v0.35 (Pucker *et al*. 2020) to identify candidate genes of the flavonoid pathway (Supplementary Methods S5). To identify MYB transcription factors, we searched the predicted protein sequences of the *A. hypochondriacus* genome annotation v2.2 for matches to the MYB DNA-binding domain HMM profile (Pfam ID: PF00249, downloaded 11.01.2022) using HMMer v3.3.2 (Eddy 2009) and used the identified proteins for phylogenetic analysis (Supplementary Methods S5). We quantified gene expression in different tissues using kallisto v0.48.0 (Bray *et al*. 2016) with sequence based bias correction and the previously described short-read datasets (Table S1).

### Mapping betalain related quantitative trait loci

To map the genetic basis of tissue color differences in amaranth, we created two distinct mapping populations. For the color regulator bulk segregant analysis (BSA), we used an F_2_ mapping population created from a cross of PI 686465, which represents an F_1_ hybrid of PI 568125 and PI 568179 (Brenner 2019), with PI 538323. For the biosynthesis BSA, the F_2_ mapping population was created from a cross between PI 686465 and PI 576485. Because PI 686465 and PI 538323 were not homozygous lines, the mapping populations might carry multiple haplotypes for functional loci. Plants were grown well-watered at 60 % air humidity under controlled short-day conditions with night and day temperatures set to 19*°C* and 24*°C*, respectively. For the regulator BSA, plants were bulked (69 plants per bulk) based on flower color, however all plants produced red leaves. For the biosynthesis BSA, plants were bulked (80 plants per bulk) based on whole plant color and green plants never produced red coloring pigments. We extracted betalains from leaves and flowers of the three parental accessions, using a modified protocol (Chang *et al*. 2021; von Elbe 2001), and determined betalain content using photometric quantification and liquid chromatography - mass spectrometry (LC-MS) analysis (Supplementary Methods S6). We extracted bulked DNA from the pools using the DNeasy Plant Pro Kit (Qiagen, Hilden, Germany) using the manufacturers protocol. Extracted DNA was sequenced by the Cologne Center for Genomics (CCG) on an Illumina NovaSeq 6000 platform in 150 bp paired-end reads (Table S4). Sequencing quality was assessed using fastQC (Andrews 2010). We mapped reads to the *A. hypochondriacus* reference genome v2.2 using bwa-mem2 v2.2.1 (Li 2013) and followed GATK v4.1.7 “best practices” (McKenna *et al*. 2010) to call and filter SNPs. Only biallelic SNPs from the 16 largest Scaffolds were included in the analysis and sites were filtered for a maximum of 5 % missing values using vcftools v0.1.16, resulting in a total of 4,367,544 SNPs (Danecek *et al*. 2011). We predicted the effect of called SNPs on protein coding genes using SnpEff (Cingolani *et al*. 2012). We carried out BSA using the QTLseqR package (Mansfeld and Grumet 2018) and increased the genome-wide false-discovery G’ threshold to 3.

To compare gene expression between the green and red bulks of both BSAs, we newly bulked 29 plants of each bulk for the biosynthesis BSA and 24 plants from each bulk for the regulator BSA. We extracted RNA from bulked flower tissue using the Invitrogen PureLink Plant RNA Reagent (Thermo Scientific™, Thermo Fisher Scientific Inc., Massachusetts, USA) following the manufacturers protocol. Sampled flower tissue was immediately stored at −80*°C* until RNA extraction. Sequencing libraries were created using the TruSeq stranded mRNA preparation kit and sequenced by the CCG on a Illumina NovaSeq 6000 platform in stranded 100 bp paired-end reads (Table S5). We removed adapter sequences using trimmomatic v0.39 (Bolger *et al*. 2014), evaluated sequencing quality using fastQC (Andrews 2010), and quantified gene expression using kallisto (Bray *et al*. 2016). We defined a TPM threshold of 0.5 in at least one sample to define a gene as expressed.

To confirm the presence of the mutually exclusive non-synonymous and stop variants in the *AhCYP76AD2* gene of green plants from the biosynthesis BSA, we Sanger sequenced (LightRun, Eurofins Genomics) one individual with red leaves and four individuals with green leaves using primers SV4 and SV5 (Table S6). Sequencing results were mapped to the *AhCYP76AD2* genomic sequence and visualized using CLC DNA Workbench 5.6.1 (QIAGEN).

### Cloning and overexpression of AhMYB2

We cloned *AhMYB2.1* and *AhMYB2.2* from cDNA of the flower tissue RNA sample of PI 558499 also used for isoform sequencing (see above). We amplified *AhMYB2* CDS by PCR with primers SV1 and SV2 (Table S6) and assembled the final vectors, using the pGGZ001, pGGA004, pGGB003, MYB isoforms in pGGC000, pGGD001, pGGE001 and pGGF008 modules of the green gate system (Lampropoulos *et al*. 2013). The two resulting vectors were *p35S:AhMYB2.1-GFP* and *p35S:AhMYB2.2-GFP*. In addition, we used *p35S:mCherry-NLS* as control for nuclear localisation.

For hairy root transformation, we co-transformed *Agrobacterium rhizogenes* AR1193 (Stougaard *et al*. 1987) with the expression vectors and the pSoup plasmid (Hellens *et al*. 2000). We performed hairy root transformation according to Lota *et al*. (2013) with minor changes. The *A. rhizogenes* cultures for hairy root transformation were grown on plates for 2 days at 28 °C. We scraped the bacteria from the plates using sterile loops and re-suspended them in PIPES buffer (Lota *et al*. 2013), using around 1.5 plates fully covered with *A. rhizogenes* biofilm per mililiter of buffer. Seeds of PI 558499 were sterilized and germinated on solid Murashige-Skoog-medium (MS) (Murashige and Skoog 1962) with 1 % sucrose. We used a sterile 27 G needle dipped into the respective bacterial solution to wound each 3-4 day-old seedlings hypocotyl several times. *Agrobacterium*-treated plants were then grown in a growth chamber at 22 °C with a 16/8 h light/dark cycle. Hairy roots were evaluated visually for their color after 23 days. We evaluated GFP signal under the Leica MZ16 F stereo microscope equipped with a DFC420 C camera and Leica 10447407 GFP long-pass filter to validate successful transformation.

To study the sub-cellular localisation of AhMYB2, we bombarded leek epidermal cells with DNA-coated gold particles following Pietsch (2022). We coated gold particles simultaneously with *p35S:mCherry-NLS* and either *p35S:AhMYB2.1-GFP* or *p35S:AhMYB2.2-GFP* to evaluate nuclear localisation. We analyzed the bombarded leek cells after 24 to 30 hours under Leica DMRE microscope with a DFC7000 T camera using the AHF F36-504 mRFP (for mCherry) and AHF F46-002 EGFP (for GFP) filter cubes.

## Results

### Improved A. hypochondriacus reference genome

A high-quality reference genome sequence is a prerequisite for numerous types of analyses. To improve the available *A. hypochondriacus* genome assembly and remove indel errors we polished the genome using 38.8 Gb of Illumina reads (Lightfoot *et al*. 2017). After filtering of reads with ambiguous basecalls, 384,876,330 reads with length of 100 bp remained. Genome polishing resulted in a length increase of 105,049 bp (from 403,889,442 bp to 403,994,491 bp) of the polished reference genome compared to the unpolished reference genome (Lightfoot *et al*. 2017). In total, 56.91 % (229,915,110 bp) of the assembly were classified as retroelements (20.57 %), DNA transposons (4.86 %), or simple repeats and low complexity regions (3.08 %; Table S7). These elements were distributed across the genome with increased density in regions with lower gene content (Fig. 1c). We further identified and annotated regions with organellar sequence similarity (Fig. S1). These regions could have resulted from erroneous assembly or could be the result of transfer into the genome (Wang *et al*. 2018).

In order to improve the annotation with full-length transcript sequencing evidence, we performed PacBio isoform sequencing of seven different tissues of the amaranth reference accession (PI 558499, Table S1, Supplementary Results S1). The assembled Iso-Seq transcriptome showed high reported completeness, which was further increased by reference-based correction using the polished reference genome (BUSCO completeness of 90.7 % and 90.8 %, respectively (Table S8). Reference-based correction using the unpolished reference genome strongly decreased the reported BUSCO score, from 90.6 % to 74.6 % complete, instead (Table S8). For a total of 43,245 transcript sequences that were predicted to be coding before and after reference-based ‘correction’ using the unpolished genome, 35.1 % (15,194) differed in predicted ORF length after correction. While 12,289 transcripts had longer predicted ORFs without ‘correction’, 2,805 were predicted with longer ORF after ‘correction’ with the unpolished reference genome. In contrast to that, only 10.1 % of transcripts differing in predicted ORF length after correction using the polished genome.

We combined the BRAKER2 genes that were at least partially supported by sequencing evidence with the predicted coding sequence of the full-length transcripts into the new genome annotation v2.2 using TSEBRA (Gabriel *et al*. 2021). Our final genome annotation has 23,817 annotated genes of which 99.12 % locate to the first 16 Scaffolds (Table 1). In comparison to the genome annotation of *A. cruentus* (89.8 % complete) and the previously published genome annotation v2.1 (80.7 % complete), the genome annotation v2.2 shows a strong increase in completeness with a BUSCO score of 98.0 % complete (Table S8). We were able to assign genes into 5,460 of 5,783 (94.41 %) of available Mercator bins of conserved biological processes and included functional annotations for 20,946 genes (95.56 %) from at least two sources (Fig. S2). These annotations will allow the effective use of the *A. hypochondriacus* reference genome for quantitative genetic analysis including genome wide association studies and molecular work.

**Table 1.** Comparison of genome annotation statistics between the *A. cruentus* genome (Ma *et al*. 2021), the published *A. hypochondriacus* genome v2.1 (Lightfoot *et al*. 2017) and the genome annotation derived from this study (v2.2). Coding sequence (CDS) and exon statistics for both previously published genome annotations were published in Ma *et al*. (2021). We calculated the BUSCO score using annotated protein sequences with identical parameters for all three genome annotations. Only the reported percentage of complete BUSCO genes (single copy and duplicated) is displayed.

|  | *A. cruentus* | *A. hypochondriacus* v2.1 | *A. hypochondriacus* v2.2 |
| --- | --- | --- | --- |
| Protein coding genes | 25,477 | 23,883 | 23,817 |
| Total isoforms | 25,477 | 23,883 | 28,074 |
| BUSCO score | 89.8 % | 80.7 % | 98.0 % |
| Mean CDS length (bp) | 1,141 | 1,066 | 1,258 |
| Mean exons per gene | 4.86 | 4.87 | 5.39 |
| Mean exon length (bp) | 235 | 219 | 233 |

### Color pathways in the A. hypochondriacus genome

To identify candidate genes for the betalain pathway, we carried out BLAST searches using the sequences of previously characterized pathway genes of different species against our annotated *A. hypochondriacus* protein sequences. For the *CYP76AD* query genes, two main candidate genes were identified, one gene as best hit for *CYP76AD1* from *B. vulgaris* and *CYP76AD2* from *A. cruentus* and one gene as best hit for *CYP76AD5* and *CYP76AD6* from *B. vulgaris* (Table S3). For *DODA1*, two different genes were identified as candidates. The candidate gene more similar to the *B. vulgaris* gene was located next to the candidate gene for *CYP76AD1* and *CYP76AD2* on Scaffold 16 (Table S3). Since multiple betalain pathway genes underwent gene duplication and neofunctionalization (Brockington *et al*. 2015), we performed phylogenetic analysis of the multiple *CYP76AD* and *DODA* candidate genes with previously described betalain pathway genes (Table S9 and Table S10). We termed the identified *CYP76AD* genes *AhCYP76AD2* and *AhCYP76AD5* for clustering with the CYP76AD *α*-clade and *β*-clade, respectively (Fig. S3). Phylogenetic analysis of identified DODA candidate genes indicated that both belonged into the DODA *α*-clade with potential DODA activity (Fig. S4). According to their relationship to BvDODA*α*1, which is involved in betalain biosynthesis (Hatlestad *et al*. 2015), and BvDODA*α*2, which displayed only minor DODA activity *in planta* and *in vitro* (Bean *et al*. 2018), we termed the two DODA candidates AhDODA*α*1 and AhDODA*α*2. We further identified three different glucosyltransferases (*AhcDOPA5GT*, *AhBetanidin5GT* and *AhBetanidin6GT*) based on the published sequences from *A. hypochondriacus* and *Amaranthus tricolor* (Table S3). We found strong expression of *AhCYP76AD2* and *AhDODAα1* especially in the red flower of PI 558499, as well as in root and stem tissue (Fig. S5). The expression patterns of glucosyltransferase genes differed between tissues. While *cDOPA5GT* showed constitutive expression in all assessed tissues, *AhBetanidin5GT* and *AhBetanidin6GT* were most strongly expressed in flower and leaf, respectively (Fig. S5). We identified candidate genes for all main betalain pathway genes in the genome annotation and annotated their genomic location.

The second major coloring pigment pathway we annotated was the flavonoid biosynthesis pathway that in other species includes the anthocyanin production. We identified flavonoid pathway genes using KIPEs (Pucker *et al*. 2020) and found copies of most flavonoid pathway genes in amaranth (Table S11). However, no candidates for *CHI2*, *F3’5’H*, *FNS1* and *ANS* could be inferred. Since the *ANR* candidate gene had low sequence similarity to known homologs and missed conserved residues, we defined the gene as missing in amaranth (Table S11). Classification into metabolic pathways using Mercator supported all genes identified by KIPEs, except for additional *F3’H* candidates, and similarly inferred *F3’5’H*, *ANS* and *ANR* to be missing. We identified multiple gene copies displaying all conserved residues for the general phenylpropanoid pathway genes *PAL*, *4CL*, and *C4H* as well as for *F3’H*. We found the general phenylpropanoid pathway genes to be strongly expressed in most tissues (Fig. S6). However the early flavonoid pathway gene *CHS* was mainly expressed in flower and water-stressed tissue (Fig. S6), possibly limiting the production of later flavonoid compounds in other tissues.

### Location and conservation of the color regulating MYB transcription factor family

MYB transcription factors are involved in the regulation of key plant processes, including anthocyanin and betalain pigment biosynthesis (Dubos *et al*. 2010; Hatlestad *et al*. 2015; Lloyd *et al*. 2017; Liu *et al*. 2015). To identify potential regulators of the pigmentation pathways, we annotated MYB transcription factor genes in the genome of amaranth. In total, we identified 82 R2R3 MYB candidate genes (92 total isoforms), three 3R MYB candidate genes (three isoforms) and one 4R MYB candidate gene in the genome of amaranth (two isoforms; Table S12). Similar to *A. thaliana* and rice (Katiyar *et al*. 2012), most R2R3 MYB genes featured two (16 genes and isoforms) or three (60 genes and isoforms) exons (Table S12). For instance, AHp022773 (*AhMYB2*) showed a conserved gene structure with three exons but two different isoforms (*AhMYB2.1* and *AhMYB2.2*) lead to different proteins and could add functional variation.

Out of the 86 identified MYB genes (R2R3, 3R and 4R), 67 could be assigned to established subgroups using phylogenetic analysis (Fig. 2; Kranz *et al*. 1998; Stracke *et al*. 2001). The identified 3R and 4R MYB genes clustered together with the 3R and 4R MYB genes of *A. thaliana* and *B. vulgaris*, respectively (Fig. 2). The conserved functions of subgroups across distant taxa enabled us to functionally assign MYBs to processes like stress responses, developmental processes or metabolite biosynthesis (Table S12). No amaranth MYBs were classified into the subgroups S6 (anthocyanin biosynthesis), S12 (glucosinolate biosynthesis), S15 (epidermal cell development) and S19 (stamen development; Fig. 2). Multiple MYB subgroups are involved in the regulation of color pathways. We classified four amaranth MYBs (AHp000789, AHp002277, AHp007908 and AHp009452) into subgroup S4, involved with transcriptional repression of different branches of the phenylpropanoid pathway (Wang *et al*. 2020). The amaranth MYBs AHp002686 and AHp017689 were classified into the early flavonoid pathway regulator subgroup S7. AHp018270 and the seed color regulation candidate AmMYBl1 (Stetter *et al*. 2020) were grouped into subgroup S5 involved with proanthocyanidin regulation. No amaranth or *B. vulgaris* MYBs were classified into the anthocyanin biosynthesis subgroup S6, but the betalain regulator BvMYB1 (Bv_jkkr; Hatlestad *et al*. 2015) was grouped together with three amaranth MYBs (AHp022773, AHp016530 and AHp016531) next to *A. thaliana* subgroup S6. The amaranth MYBs were assigned to the BvMYB1-clade, potentially associated with the regulation of betalain biosynthesis (Hatlestad *et al*. 2015). As candidate betalain pathway regulators, we named the transcription factors *AhMYB2* (AHp022773), *AhMYB3* (AHp016530) and *AhMYB4* (AHp016531). While we observed only low expression of *AhMYB3* and *AhMYB4* across all tissues, the two isoforms of *AhMYB2* were strongly expressed in flower tissue (Fig. S7).

**Figure 2.**
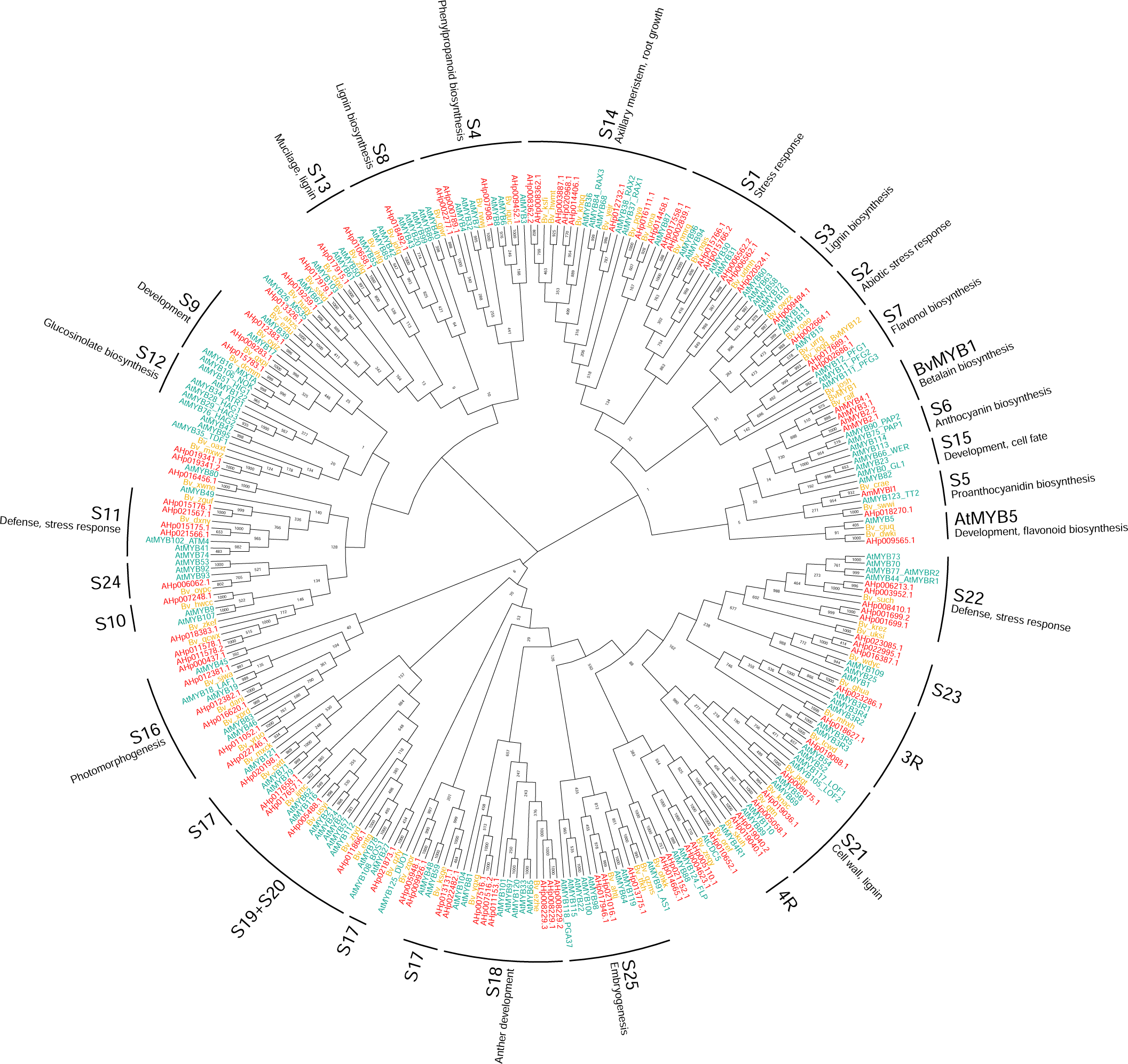
Neighbour-joining phylogenetic tree based on multiple sequence alignment of the R2R3, 3R and 4R MYB protein sequences of *A. hypochondriacus* (red), *A. thaliana* (green; Stracke *et al*. 2001) and *B. vulgaris* (yellow; Stracke *et al*. 2014). Bootstrap values from 1000 replicates, MYB subgroup assignments and functional classifications for subgroups were annotated in the phylogeny.

Different flavonoid pathway regulating MYB proteins require the formation of the MBW protein complex with two additional transcription factors (bHLH and WDR) to carry out their function (Li 2014). The interaction between MYB proteins and the bHLH interaction partners for flavonoid pathway regulation is mediated by a conserved interaction motif in the R3 domain (Zimmermann *et al*. 2004; Millard *et al*. 2019). In contrast to anthocyanin regulators, the betalain regulating *BvMYB1* has lost this interaction motive and functions without known interaction partners (Hatlestad *et al*. 2015). We therefore evaluated the presence of the conserved bHLH-interaction motif ([D/E]Lx_2_[R/K]x_3_Lx_6_Lx_3_R) in all identified amaranth MYBs. Similar to inferences in other plant species (Zimmermann *et al*. 2004; Rajput *et al*. 2022), we detected the motif in the MYB domain in two out of four genes of S4 (AHp002277 and Ahp009452), and both genes of S5 (*AmMYBl1* and AHp018270), indicating potential interaction with bHLH proteins and transcriptional regulation through the MBW complex. A slightly altered bHLH-interaction motif in which a conserved leucine residue was replaced by the biochemically similar isoleucine ([D/E]Lx_2_[R/K]x_3_Lx_6_Ix_3_R) was found in two *A. hypochondriacus* MYBs, AHp009565 from the AtMYB5-clade and AhMYB2.1 from the BvMYB1-clade. The motif differentiates AhMYB2.1 from other BvMYB1-clade proteins, in which the first residues of the motif are not conserved.

### Mapping pigment biosynthesis pathway genes

Identifying the evolutionary changes that impact color variation within populations is essential to understand the mutual exclusive nature of anthocyanins and betalains. To study the genetic basis of tissue color and its regulation in amaranth, we conducted two bulk segregant analyses (BSA) from different initial crosses (Fig. 3a). We quantified tissue color in the parental accessions and found high betacyanin content (e.g.,amaranthin) in red samples, whereas betaxanthin content was low in all assessed tissues and accessions (Figs 3, S8). In the biosynthesis BSA, 80 green (no red pigmentation in any visible tissue) and 80 red (red in all visible organs) plants were sequenced in two pools (Fig 3a). A comparison of allele frequencies between bulks identified a large QTL on Scaffold 16, from 1.1 mb to 14.4 mb (from 1,080,628 bp to 14,358,251 bp; Fig. 4a,b). The two betalain pathway genes *AhCYP76AD2* and *AhDODAα1* were located on the QTL between the two peaks and represent likely candidate genes for loss of coloration (Fig. 4b). We quantified gene expression from bulked flower tissue for the green and red bulks of both BSAs using RNA sequencing. However, gene expression levels for *AhCYP76AD2* and *AhDODAα1* were similar between the red and green bulk of the biosynthesis BSA (Fig. S9).

**Figure 3.**
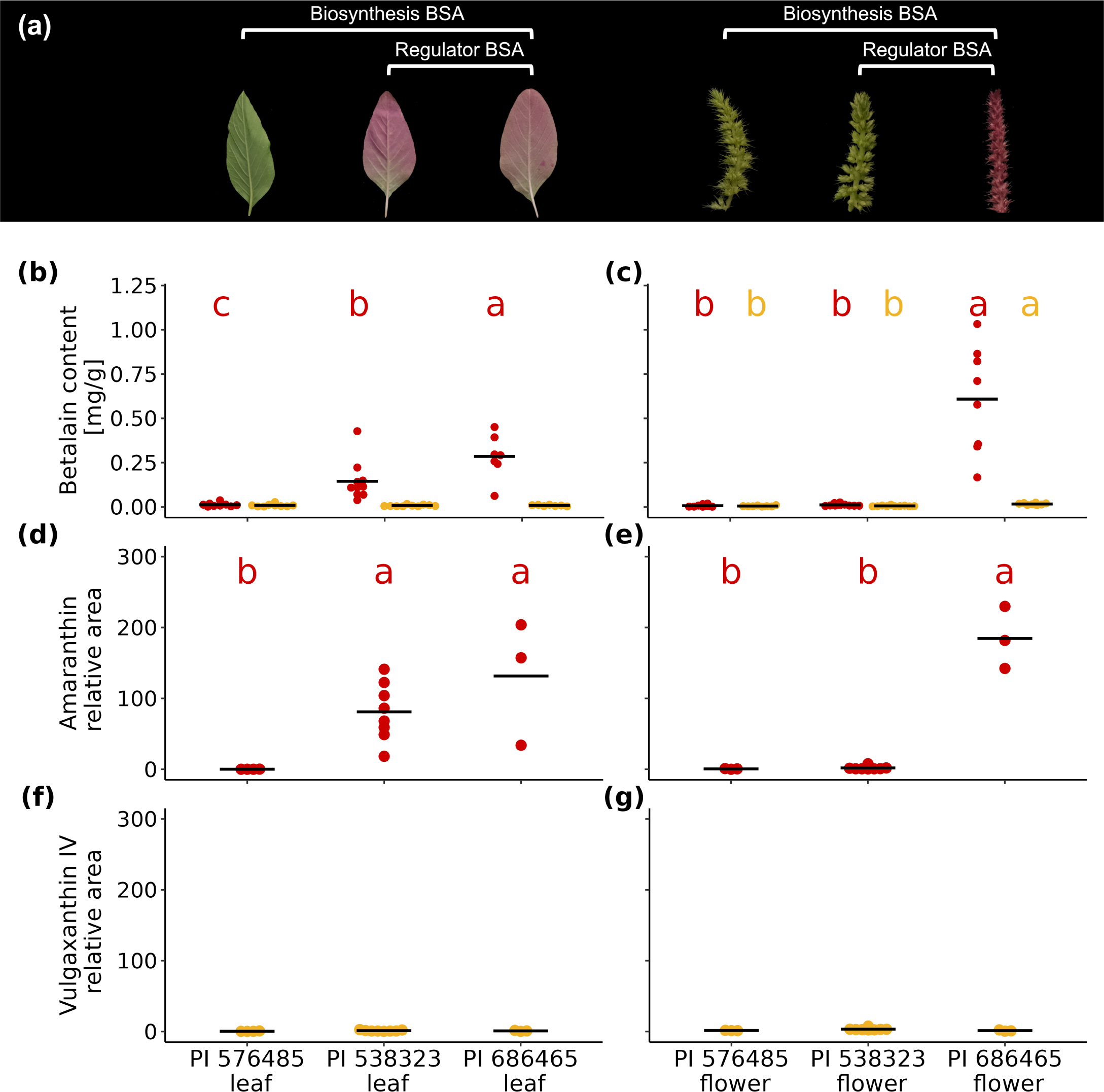
Quantification of betalain content for parental accessions used in mapping. (a) Representative leaf and flower images for parent plants, with PI 576485 and PI 686465 generating the biosynthesis BSA population and PI 538323 and PI 686465 generating the regulator BSA population. (b-c) Photometric quantification of betacyanin (red) and betaxanthin (yellow) content of (b) leaf and (c) flower tissue (mg betalains per g fresh weight), with black bars denoting mean values. (d-e) LC-MS quantification of amaranthin (betacyanin) content in (d) leaf and (e) flower tissue. (f-g) LC-MS quantification of vulgaxanthin IV (betaxanthin) content in (f) leaf and (g) flower tissue. Statistical analysis was conducted using ANOVA and significant differences in betaxanthin or betacyanin content between accessions within tissues were denoted using compact letter display. There were no significant differences in total betaxanthin content between accessions in leaf and no significant differences in vulgaxanthin IV content between accessions in leaf or flower.

**Figure 4.**
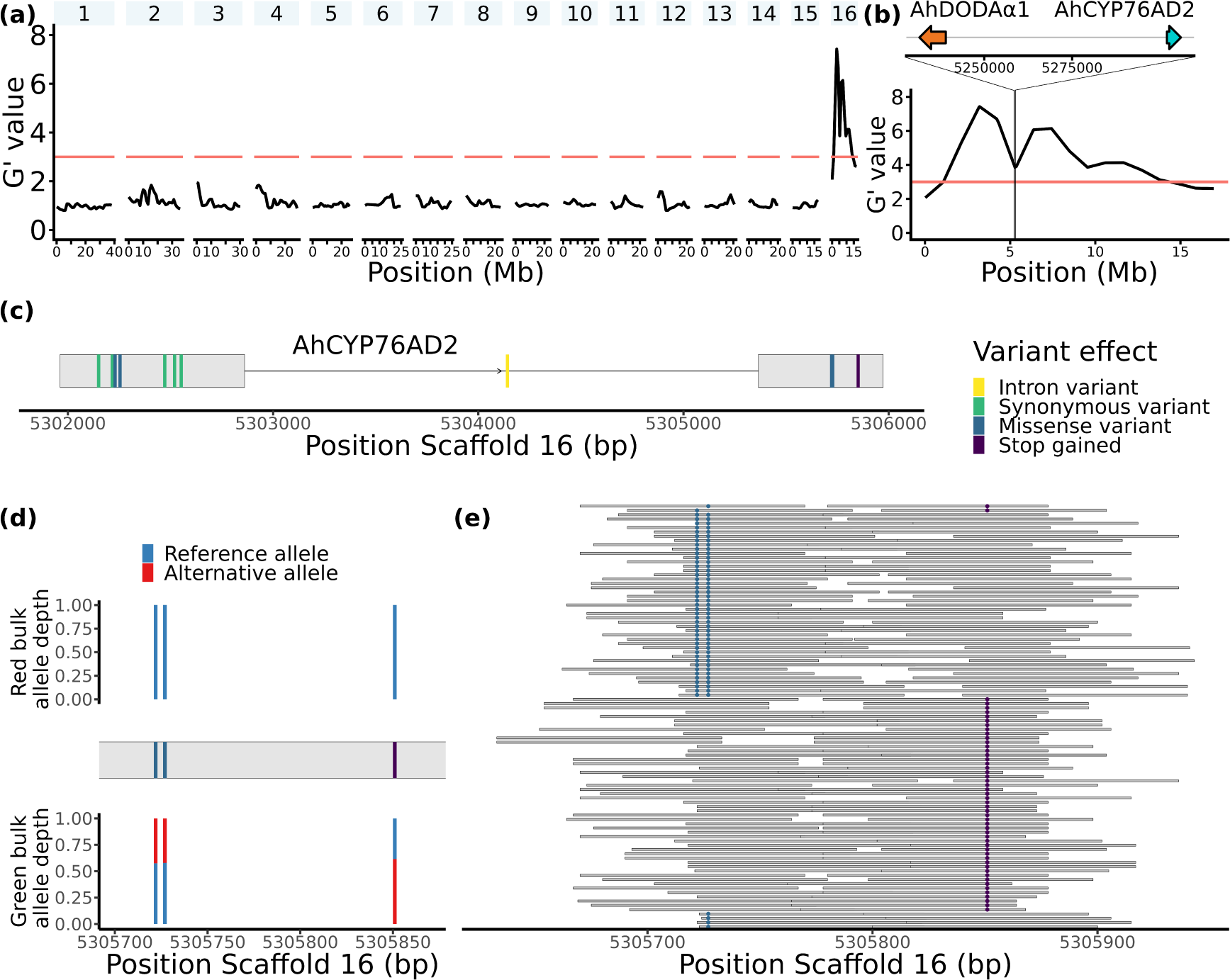
Biosynthesis mapping identifies functional variants in *AhCYP76AD2*. (a-b) Biosynthesis bulk segregant analysis (BSA) results (a) genome-wide and (b) on Scaffold 16 show QTL region on Scaffold 16. Marked by lines are the location of *AhDODAα1* (cyan) and *AhCYP76AD2* (orange) on the QTL. (c) Variants segregating in whole-genome sequencing data from red and green bulk are displayed with assigned effect of the alternative allele at their position in the exon/intron structure of *AhCYP76AD2*. (d) Position of T1258A and A1263T (missense, dark blue) and C1387T (stop, purple) variants in the second exon of *AhCYP76AD2* and their relative allele depth in the red and green bulk of the biosynthesis BSA. (e) All RNA sequencing read pairs from green bulk flower tissue of the biosynthesis BSA covering both at least one missense and the stop variant, with marked occurrences of T1258A, A1263T and C1387T alternative alleles.

To identify functional variants which could lead to color loss, we used SnpEff (Cingolani *et al*. 2012) to annotate the effect of each variant which segregated in at least one of the two bulks. We searched betalain pathway genes for segregating ‘high impact’ variants and identified missense variants and a stop-gain variant in the *AhCYP76AD2* gene (Fig. 4c) that we previously annotated as the most likely functional homolog for this enzyme. While most annotated variants occurred at similar frequencies in both bulks, two missense variants (F to I, T1258A and L to F, A1263T) and the stop gain variant at C1387T, all located on exon 2, differed strongly in relative allele depth between the two bulks (Figs 4d, S10). The SNP C1387T was monomorph for the reference allele in the red bulk of the biosynthesis BSA (allele depth of 0/24), but the alternative allele occurred at intermediate frequency in the green bulk (allele depth of 16/26). Similarly, the reference alleles of T1258A and A1263T were monomorph in the red bulk but the alternative alleles were found at intermediate frequency (relative allele depth 17/40 and 16/38, respectively) in the green bulk. The QTL included two peaks at 3.2 mb (3,205,818 bp) and 7.4 mb (7,425,588 bp) with a decreased G’ value in between, which nonetheless did not go below the significance threshold (Fig. 4b). A potential explanation might be the presence of multiple causal variants in the bulks, given their multi-parent origin (see Methods).

Since the RNA-seq data had higher coverage of *AhCYP76AD* than the whole genome sequencing data, we investigated the presence and absence of the three variants in RNA-seq data from the bulked floral tissue of additional red and green colored pools from the same cross. To resolve *AhCYP76AD2* haplotypes, we considered only read pairs which covered at least one of the non-synonymous variants and the stop variant C1387T. Inspection of read pairs originating from the green bulk revealed that either the alternative alleles of the non-synonymous variants (T1258A and A1263T) or the alternative allele of the stop variant C1387T were found in all read pairs from the green bulk (Fig. 4e). The presence of alternative alleles at T1258A and A1263T and at C1387T was almost mutually exclusive in green plants, with only two reads containing alternative alleles at C1387T and one of the missense variants. In contrast to this, less than half of all read pairs covering the *AhCYP76AD2* variants in the red bulk featured either alternative allele at T1258A, A1263T and C1387T (Fig. S11), consistent with heterozygous allele frequencies at these variants. Sanger sequencing of four individuals from the green bulk and one individual from the red bulk confirmed the presence of mutually exclusive alleles in the sampled green bulk (Fig. S12). Together, this suggests that we mapped two functional haplotypes that both disrupt the first enzyme in the betalain pathway, leading to loss of red betacyanins in the whole plant.

### Flower-specific regulation of betalain enzymes by AhMYB2

We further intended to identify tissue-specific regulators of betalain pigmentation in amaranth. Therefore, we conducted a second BSA from a different initial cross and pooled 69 plants based on red or green flower color (Fig. 3a). All plants from both pools had red leaves and hence could produce betalains (Fig. 3). In the regulator BSA, we identified a QTL for flower color at the beginning of Scaffold 16 up to 5.2 mb (1-5,244,109 bp; Fig. 5a,b). We found the newly annotated betalain pathway regulator candidate *AhMYB2* to be located at the center of the QTL (Fig. 5b). While *AhCYP76AD2* and *AhDODAα1* were also covered by the QTL, we did not find large effect mutations differing between pools and both pools did synthesise red betalains in leaves (Fig. 3). We did not identify any ‘high impact’ mutations in *AhMYB2*, but whole RNA sequencing of flower tissue from pools of diverging flower color individuals revealed a strongly reduced gene expression of both annotated *AhMYB2* isoforms in green flowers compared to red flowers (Fig. 5c). This indicates that the loss of *AhMYB2* expression is associated with the loss of flower color. Comparison of betalain pathway enzyme gene expression revealed reduced expression levels of *AhCYP76AD2* and *AhDODAα1* in green flowers (Figs 5c, S9). Notably, we did not detect expression of *AhCYP76AD5* and downstream betalain pathway gene expression did not differ between red and green flowers, indicating that reduced *AhCYP76AD2* and *AhDODAα1* gene expression caused betalain loss in green flowers (Figs 5c, 1b). Given the difference in flower color between pools in the regulator BSA (Fig. 3), strong expression of *AhMYB2* only in flower tissue (Fig. S7) and the correlation of its expression with that of two betalain pathway genes, *AhMYB2* likely represents a flower-specific regulator of the betalain pathway (Fig. 5c).

**Figure 5.**
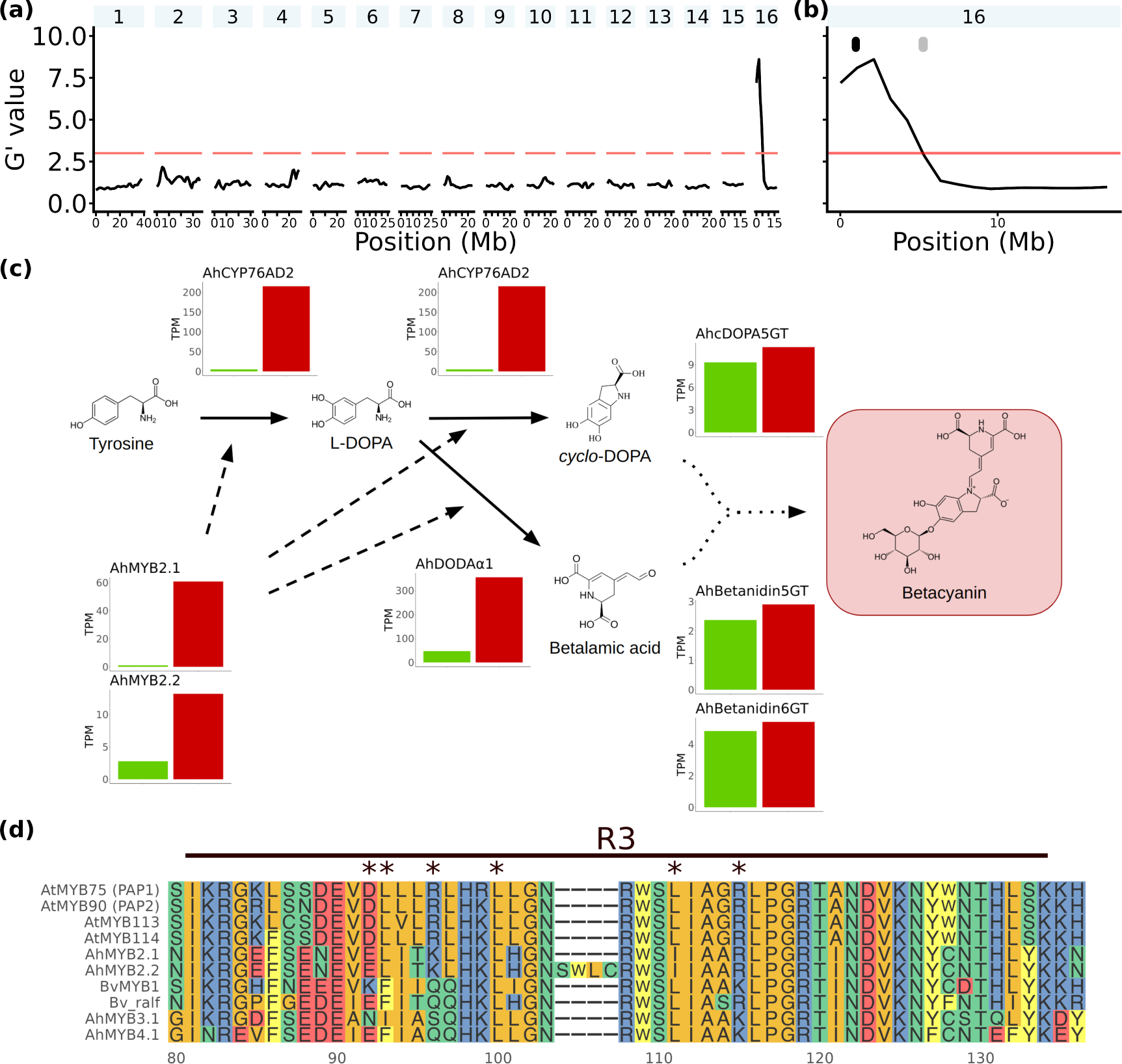
Regulator mapping identifies the flower color regulator *AhMYB2*. (a-b) Regulator BSA results (a) genome-wide and (b) for Scaffold 16 only show the identified QTL region (1-5,244,109 bp) on Scaffold 16. The QTL region includes the betalain pathway genes *AhCYP76AD2* and *AhDODAα1* (grey) as well as the regulator *AhMYB2* (black). (c) Proposed model of regulation of early betalain pathway gene expression by *AhMYB2* in a simplified pathway depiction alongside relative gene expression [TPM] in flower tissue for betalain pathway genes in green and red flower bulk of the regulator BSA. Multiple enzymatic reactions and paths leading to red betacyanin production were simplified with dotted arrows. The structure of betanin was displayed as representative betacyanin. The potential betalain pathway genes *AhMYB3*, *AhMYB4* and *AhCYP76AD5* were not expressed. (d) Alignment of the R3 domain of *A. thaliana* anthocyanin regulators with BvMYB1-clade proteins. Residues of the bHLH-interaction motif as defined by Zimmermann *et al*. (2004) ([D/E]Lx_2_[R/K]x_3_Lx_6_Lx_3_R) were marked with asterisks. Residue 111 of the alignment constitutes the imperfectly conserved interaction motif residue in AhMYB2.1.

The protein sequences of the AhMYB2 isoforms AhMYB2.1 and AhMYB2.2 differed only by four additional residues in AhMYB2.2, stemming from an earlier start of exon 3 (Fig. 5d). The additional residues in AhMYB2.2 disrupt the conserved R3 domain and the bHLH-interaction motif, which might impair the function of this isoform. To functionally validate *AhMYB2* as regulator of the betalain pathway, we overexpressed both isoforms in transgenic hairy roots in amaranth and investigated their cellular localisation in leek (Fig. 6a-f). As expected for transcription factors, both isoforms were localised in the nucleus (Fig. 6c,f). *AhMYB2.1* overexpression resulted in red coloration of emerging roots, validating the suggested involvement in betalain pathway regulation (Fig. 6a,b). In contrast, *AhMYB2.2* overexpression resulted only in the formation of white roots (Fig. 6d,e), indicating that the insertion in the conserved MYB domain does indeed disrupt its function and further validating the function of *AhMYB2.1*. In AhMYB2.1, we identified a slightly altered version of the bHLH-interaction motif compared to that found in anthocyanin regulators, in which a conserved leucine residue was replaced by isoleucine ([D/E]Lx_2_[R/K]x_3_Lx_6_Ix_3_R; Fig. 5d). Presence of the motif differentiates AhMYB2.1 from other BvMYB1-clade proteins, in which the first residues are not fully conserved (Fig. 5d). Possible interaction of AhMYB2.1 in the MBW complex could indicate an undiscovered mechanism of betalain regulation more similar to anthocyanin regulators from other species.

**Figure 6.**
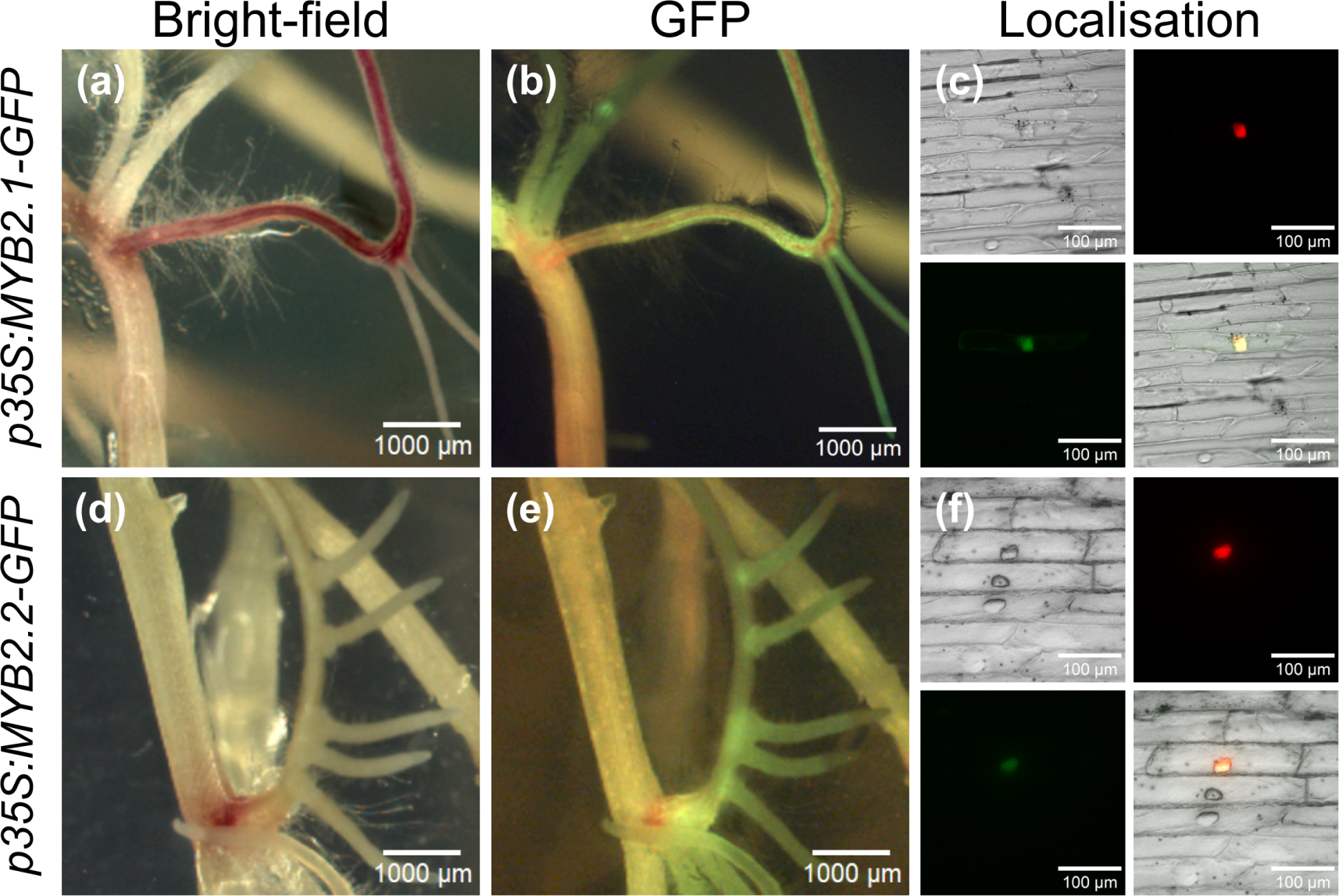
Functional validation of *AhMYB2* as betalain regulator. (a-c) Over-expression of *AhMYB2.1* in *A. hypochondriacus* hairy roots, (a) bright-field and (b) GFP-channel show GFP-tag signal and red pigmentation phenotype. (c) Nuclear localisation of *p35S:AhMYB2.1-GFP* in leek epidermins after co-bombardment with *p35S:NLS-mCherry*. (d-f) Over-expression of *AhMYB2.2* in *A. hypochondriacus* hairy roots, (d) bright-field and (e) GFP-channel show GFP-tag signal but no red coloration. (f) Nuclear localisation of *p35S:AhMYB2.2-GFP* in leek epidermins after co-bombardment with *p35S:NLS-mCherry*. (c, f) Subplots show bright-field (top left), mCherry (top right), GFP (bottom left) and merged (bottom right) channels.

## Discussion

### High quality reference genome annotation for functional genomics

We improved the available reference genome of *A. hypochondriacus* (Lightfoot *et al*. 2017) by genome polishing and subsequent genome annotation. While the *A. hypochondriacus* assembly was highly contiguous with 16 chromosome-scale Scaffolds, even single nucleotide insertions or deletions in coding sequences could have a large impact on gene prediction through frame shifts. The increased completeness of our genome annotation v2.2 compared to the previous annotation likely stems from the removal of a large number of small insertions and deletions in the previous assembly, which impaired protein prediction. Higher similarity of long-read transcript sequences to the polished reference genome and higher overall annotation completeness after genome polishing demonstrates the successful removal of small errors in the assembly (Table S8). By using the polished genome for genome annotation and adding additional full-length transcript sequencing data, we were able to produce the most complete genome annotation of amaranth to date (Clouse *et al*. 2016; Lightfoot *et al*. 2017; Ma *et al*. 2021; Wang *et al*. 2023) and included the annotation of multiple isoforms for 3,484 genes (Table 1).

Alternative splicing is involved in regulatory processes and greatly expands on the coding capacity of eukaryotes through the production of alternative isoforms (Wang *et al*. 2016; Kornblihtt *et al*. 2013). Our full-length transcript sequencing data allowed for accurate annotation of different splice variants in grain amaranth. These alternative splice variants will enable future functional studies on isoform-specific gene expression, as shown for the newly annotated *AhMYB2* gene for which we found one stronger expressed, and functional splice variant (Figs 5c, 6). This splicing variation potentially allows for additional functions or differential regulation of the gene in other organs (Huang *et al*. 2022; Kornblihtt *et al*. 2013). While we were able to annotate multiple isoforms for over three thousand genes, additional isoform sequencing in further tissues and accessions will be needed to represent the complete splicing variation in the species.

The reported completeness of the new genome annotation is comparable in accuracy to well annotated model plant genomes (Dohm *et al*. 2014). It represents a valuable resource that will allow more accurate studies of regulatory processes and support functional genetics and future breeding efforts in amaranth.

### Conservation of the flavonoid and betalain pigment pathways in amaranth, but loss of anthocyanin pathway genes

While betalains and anthocyanins appear to be mutually exclusive, the flavonoid pathway upstream of anthocyanin production is reported to be intact in betalain producing species and loss or downregulation of only specific anthocyanin genes appears to have led to the loss of anthocyanins in the Caryophyllales (Sakuta *et al*. 2021; Pucker *et al*. 2023). To study the conservation of the flavonoid pathway in amaranth, we identified and annotated genes of the general phenylpropanoid and flavonoid pathway using our improved genome annotation. We found no close homolog of *FNS1*, corresponding to previous reports of *FNS1* homologs being frequently absent outside of Apiaceae species (Pucker *et al*. 2020; Gebhardt *et al*. 2007). The amaranth genome did not contain a *F3’5’H* homolog, which is absent in various plant species (Table S11; Wessinger and Rausher 2012). Still, we found the flavonoid pathway to be generally intact in amaranth with loss of anthocyanidin synthase (*ANS*) and key residues in anthocyanidin reductase (*ANR*) supporting the complete loss of anthocyanin production in amaranth.

While the genes of the betalain pathway have been identified in different amaranth species in previous studies (Chang *et al*. 2021; Zheng *et al*. 2016; Casique-Arroyo *et al*. 2014), no comprehensive annotation of the pathway in *A. hypochondriacus* was available that would enable candidate gene association in mapping studies (Table S3). The characterization of functional betalain genes was complicated by multiple ancient gene duplications in the Caryophyllales, which gave rise to specific CYP76AD and DODA lineages involved in betalain production (Brockington *et al*. 2015; Sheehan *et al*. 2020). Phylogenetic analysis showed that *AhCYP76AD2* and *AhDODAα1* belong to their respective *α*-clades and are likely functioning in betalain biosynthesis (Figs S3, S4; Sunnadeniya *et al*. 2016; Brockington *et al*. 2015). In contrast to *AhCYP76AD2*, *AhCYP76AD5* from the CYP76AD *β*-clade is likely only capable to catalyze the conversion of L-DOPA to cyclo-DOPA (Brockington *et al*. 2015). Similar to previous reports, we found *AhCYP76AD2* and *AhDODAα1* to be co-localized on a conserved betalain locus on Scaffold 16 (Fig. 4b; Lightfoot *et al*. 2017; Ma *et al*. 2021). The biosynthesis BSA identified this locus to be associated with red plant pigmentation (Fig. 4), further supporting the involvement of the locus in betacyanin biosynthesis. We identified mutually exclusive alleles in the *AhCYP76AD2* gene, associated with complete loss of red tissue color (Fig. 4). Studies on loss of anthocyanin meditated flower color show that, while functional (loss-of-function in pathway genes) and regulatory mutations (leading to e.g. gene downregulation in specific tissues) occur at comparable frequencies, regulatory mutations leading to color loss appear to be preferentially fixed in natural populations, likely because of pleiotropic effects of pathway genes in vegetative tissue (Wessinger and Rausher 2012; Gates *et al*. 2018). In our population, betalain pathway genes showed even higher expression in the green bulk compared to the red bulk (Fig. S9). Since betalain genes were still expressed in green tissue, causal mutations for the loss of tissue color likely led to a knock-out of *AhCYP76AD2* (Fig. 4). While the knock-out allele at C1387T shows high evidence to lead to a loss of *AhCYP76AD2* function and the loss of betacyanin biosynthesis, the missense alleles at T1258A and A1263T might lead to the loss of function by changed protein folding or through the replacement of functionally important residues (Fig. 4c,d). The presence of independent functional mutations and the complete lack of red color might indicate dispensability of betacyanin production, at least in individual populations. Studies on the fitness effect of betacyanin loss and the prevalence of knock-out mutations and their possible pleiotropic effects on plant fitness in natural amaranth populations are currently lacking.

We found conserved patterns in the betalain and flavonoid pathways through pathway gene annotation in the improved gene atlas, but the degradation of key anthocyanin enzymes. However, the loss of anthocyanin genes does not have to be causal for mutual exclusiveness, since e.g., previous downregulation or replacement of anthocyanin genes would remove selective constraints to maintain the genes. Since other flavonoids can still be synthesised in betalain producing species (Iwashina 2015), the emergence of betalain biosynthesis alone is likely not sufficient to explain total replacement of anthocyanins and the observed mutual exclusiveness of the pigments across the large phylogenetic range of the Caryophyllales. To understand the subsequent order of the rise of mutual exclusiveness between betalains and anthocyanins, it might be important to understand the regulation of both pathways in the same study system.

### AhMYB2 is a novel regulator of the betalain pathway

Despite the characterisation of the betalain biosynthesis pathway in recent years (Hatlestad *et al*. 2012; Vogt *et al*. 1999; Christinet *et al*. 2004) and the identification of few regulators (Xie *et al*. 2023; Zhang *et al*. 2021; Zeng *et al*. 2023; Hatlestad *et al*. 2015), knowledge about tissue-specific regulation of the pathway is limited. We identified the novel amaranth flower color regulator *AhMYB2* in our regulator mapping population (Fig. 5). Strongly reduced gene expression of *AhMYB2*, as well as of the betalain pathway genes *AhCYP76AD2* and *AhDODAα1* in green flowered plants of the regulator BSA (Fig. 5c) leads us to propose *AhMYB2* as transcriptional regulator of the early betalain pathway. While we annotated two different *AhMYB2* isoforms and both showed reduced expression in green flower in the regulator BSA, an insertion in the conserved MYB domain leaves only *AhMYB2.1* capable of inducing betalain accumulation (Figs 5d, 6). Flavonoid pathway regulating MYB transcription factors commonly control tissue-specific pigment accumulation and frequently determine specificity of the MBW complex (Rajput *et al*. 2022; Lin-Wang *et al*. 2010; Xu *et al*. 2015). While *BvMYB1* was shown to regulate the betalain pathway in *B. vulgaris* leaves (Hatlestad *et al*. 2015) its ortholog *AhMYB2* regulates betalain biosynthesis in flower, instead. Similar to the tissue-specific regulation by flavonoid pathway MYBs, other BvMYB1-clade proteins might represent regulators in other tissues. While we did not observe high expression for *AhMYB3* and *AhMYB4* in any sampled tissue (Fig. S7), the genes might be active in different conditions or genotypes and should be targets of characterization in future studies. The identification of *AhMYB2* as flower-specific color regulator improves the understanding of the tissue-specific regulation of the early betalain pigment pathway.

The flavonoid pathway in plants is regulated by conserved subgroups of MYB transcription factors (Gonzalez *et al*. 2008; Lloyd *et al*. 2017; Liu *et al*. 2015). We found the conservation of phenylpropanoid pathway (S4), early flavonoid pathway (S7) and proanthocyanidin (S5 and AtMYB5-like) MYB regulators in amaranth, in accordance with the conservation of the flavonoid pathway enzymes (Fig. 2, Table S12). Clustering of AhMYB2 with the betalain regulator BvMYB1 next to anthocyanin MYBs (S6) supports the proposed neofunctionalization of former anthocyanin regulators for betalain pathway regulation (Figs 2, 5, Table S12; Hatlestad *et al*. 2015; Timoneda *et al*. 2019). While MBW complex interaction is conserved in S6 anthocyanin regulators across species (Zimmermann *et al*. 2004; Xu *et al*. 2015), the ability to interact with bHLH proteins in the MBW complex might have been lost in the betalain regulator BvMYB1 (Hatlestad *et al*. 2015). In contrast to other *A. hypochondriacus* and *B. vulgaris* BvMYB1-clade MYBs (Hatlestad *et al*. 2015), AhMYB2.1 includes an altered version of the conserved bHLH-interaction motif in the R3 domain (Fig. 5d), which it shares with the amaranth MYB AHp009565. The homolog of AHp009565 in *A. thaliana*, AtMYB5, contains an canonical interaction motif (Zimmermann *et al*. 2004), suggesting that the altered bHLH-interaction motif might still enable the amaranth MYBs to interact with either known or novel bHLH proteins. Through possible MBW complex formation, AhMYB2 might regulate the betalain pathway via a currently unknown mechanism similar to known anthocyanin regulators.

### The rise of mutually exclusive pigment pathways

Within the Caryophyllales, betalain producing clades are intermixed with anthocyanin producing lineages, leading recent studies to propose multiple evolutionary origins of betalains (Sheehan *et al*. 2020; Timoneda *et al*. 2019). Despite this polyphyletic origin of betalain production, the biosynthesis of betalains and anthocyanins appears to be mutual exclusive in all plants (Clement and Mabry 1996). A deregulated form of the arogenate dehydrogenase gene has been suggested to have enabled the shift from anthocyanins to betalains, by increasing carbon flux of the common substrate arogenate towards the betalain precursor tyrosine and thereby decreasing the availability of phenylalanine for flavonoid (including anthocyanin) production (Lopez-Nieves *et al*. 2018; Timoneda *et al*. 2019). Yet, changes in substrate availability cannot sufficiently explain the mutual exclusiveness of the two pigments, since many other flavonoids are still readily synthesised in betalain producing species (Iwashina 2015). Both, expression differences in anthocyanin biosynthesis genes and loss of anthocyanin transport and flavonoid pathway genes contribute to prevent anthocyanin production in betalain species (Shimada *et al*. 2007; Pucker *et al*. 2023). Such regulatory or functional changes of anthocyanin genes can explain how simultaneous production of both betalains and anthocyanins is never observed, but not the evolutionary reason for the alternation. Step-wise models of the shift from anthocyanin to betalain biosynthesis commonly include intermediate phases with both anthocyanin and betalain production and only later loss of the functionally equivalent anthocyanins (Timoneda *et al*. 2019). A potential key to explain the mutual exclusiveness might be to focus instead on the common components utilized by both pathways, which includes the MYB regulators (Hatlestad *et al*. 2015). Previous studies found orthologs of anthocyanin MYBs in betalain producing species unable to activate expression of genes required for anthocyanin biosynthesis (Shimada *et al*. 2007; Sakuta *et al*. 2021). The co-option of MBW complex interacting anthocyanin MYBs into betalain regulation, followed by loss of key residues for bHLH-interaction could have relieved anthocyanin genes of their key transcriptional regulators (Timoneda *et al*. 2019; Hatlestad *et al*. 2015; Sakuta *et al*. 2021). In this context, the altered bHLH-interaction motif identified in AhMYB2 could represent an interesting intermediate step in the proposed regulator shift. Further characterization of *AhMYB2* and other betalain regulators might help to understand the co-option of former anthocyanin regulators into the regulation of the novel betalain pigment pathway, the molecular prerequisites for this change and the mutual exclusiveness of the two pathways.

## Acknowledgments

We thank David M. Brenner from Iowa State University and a Curator in the US National Plant Germplasm System for providing F_2_ mapping populations for pigmentation gene identification. We thank Roswitha Lentz for help with DNA & RNA extraction and Kerstin Becker and Karl Koehrer from the Genomics & Transcriptomics Labor (BMFZ) at HHU for help with the PacBio sequencing. Analysis was conducted on the Regional Computing Center of the University of Cologne (RRZK) High Performance Computing (HPC) system CHEOPS (Funding number: INST 216/512/1FUGG). We thank Jessica Pietsch, Hanna Bechtel and Swen Schellmann for the help with the bombardment and advice on the microscopy and Jathish Ponnu for providing the *p35S:mCherry-NLS* vector. We acknowledge funding by the Deutsche Forschungsgemeinschaft (DFG, German Research Foundation) under Germanýs Excellence Strategy – EXC-2048/1 – Project ID 390686111 and STE 2654/4. TSW was supported by the University of Cologne Forum for Plant Biology and Politics of Nutrition (BiPoN).

## Data availability

All sequencing data generated in this study are deposited under BioProject ID PRJEB65083. Code used for the study is available at https://github.com/T-Winkler/Ahyp_v2_2_publication.git. The amaranth genome and annotation v2.2 can be accessed through CoGe under Genome id66602 https://genomevolution.org/coge/ GenomeInfo.pl?gid=66602. All files, including functional annotations, can be accessed at https://doi.org/10.6084/ m9.figshare.23946765.

## Competing interests

None declared.

## Author contributions

TSW and MGS planned and designed the study. TSW and MGS performed experiments and analysed the data. SKV amplified and sequenced *AhCYP76AD2* haplotypes. SKV cloned and overexpressed AhMYB2. NDR and SM performed LC-MS analysis. TSW, NDR and SKV prepared the figures. TSW and MGS wrote the manuscript. All authors read and approved the manuscript.

## Supporting Information

### Supplement

### Supplementary Methods

#### Methods S1: Genome polishing and masking of repetitive elements

To prepare the downloaded reference genome for genome polishing, we removed stretches of ambiguous bases, usually located at the beginning or end of scaffolds, using a custom script (https://github.com/T-Winkler/ Ahyp_v2_2_publication.git). Ambiguous bases were merged back into the genome after genome polishing. We ran two cycles of polishing using the two NextPolish core modules, k-mer score chain and k-mer count (Hu *et al*. 2020). Before each polishing module, we mapped reads using bwa-mem2 v2.2.1 (Li 2013) and processed bam files using Samtools v1.13 (Li *et al*. 2009). After genome polishing, we masked the polished genome for repetitive elements in order to prepare the computational gene prediction, using Repeatmodeler v2.0.1 (Flynn *et al*. 2020) with the *-LTRStruct* option to create a database of repetitive elements from the polished assembly. We reclassified repetitive elements using RepeatClassifier v2.0.4 (Flynn *et al*. 2020) and used the repetitive element database to annotate repeats in the polished reference genome using Repeatmasker v4.1.1 (Chen 2004) with the *-e rmblast* option. We softmasked the polished genome using bedtools v2.29.2 (Quinlan and Hall 2010) *maskfasta* with the repetitive element annotation to generate the *A. hypochondriacus* genome assembly v2.2. To identify genomic regions showing mapping bias, we mapped the above described whole-genome sequencing reads to the polished reference genome using bwa-mem2 v2.2.1 (Li 2013) and calculated read depth using Samtools v1.13 (Li *et al*. 2009). We identified organellar contamination within the assembly by mapping the *B. vulgaris* mitochondrial genome (Kubo *et al*. 2000) and the *A. hypochondriacus* chloroplast genome (Chaney *et al*. 2016) against the polished assembly using minimap2 v2.17 (Li 2018) with the *asm10* and *asm5* presets, respectively.

#### Methods S2: Assembly of full-length Iso-Seq transcripts

Circular consensus sequence (CCS) reads were processed using Isoseq3 v3.3.0 (https://github.com/PacificBiosciences/ IsoSeq; Table S2). After primer removal and demultiplexing of the CCS reads, poly-A tails were trimmed and artificial concatemers were removed using *Isoseq3 refine* to create a set of full-length non-concatemer (FLNC) reads (requiring FL reads to include a poly-A tail >= 20 bp). We combined the FLNC reads from all sequenced tissues using Samtools v1.13 (Li *et al*. 2009) *merge*. We clustered full-length non-concatemer reads of the seven different tissues into a set of high quality isoforms with predicted accuracy *≥* 0.99 using *Isoseq3 cluster*. We mapped the clustered high quality isoforms to the polished *A. hypochondriacus* reference genome using minimap2 v2.17 (Li 2018) with the *splice:hq* preset and the *–secondary=no* and *-uf* options. In order to remove redundant sequences, we collapse the mapped high quality isoforms into unique isoforms using the *collapse_ isoforms_by_sam.py* script of cDNAcupcake v28.0.0 (https://github.com/Magdoll/cDNA_Cupcake). Only mapped high quality isoforms with a coverage of *≥* 0.95, identity *≥* 0.9, an identical intron chain and maximum 3’ and 5’ end differences of 1000 bp were collapsed into unique full-length transcripts. We obtained the number of clustered high quality isoforms supporting each unique full-length transcript and filtered possible 5’ degraded transcripts with the *cDNA cupcake* scripts *get_abundance_post_collapse.py* and *filter_away_subset.py*.

We used SQANTI3 v4.2 (Tardaguila *et al*. 2018) for quality control of the unique isoforms as well as potential sequencing error correction. SQANTI3 uses the genomic positions of the mapped unique full-length transcripts to correct possible sequencing errors using the reference genome sequence (Tardaguila *et al*. 2018). We predicted the coding potential of the corrected transcript sequences as well as putative open reading frames (ORFs) in the sequences using CPC2 v1.0.1 (Kang *et al*. 2017). Transcripts classified as protein-coding were required to be labeled “coding” with positive ORF integrity. To estimate the completeness of the generated transcripts, we used BUSCO (Simão *et al*. 2015) to compare the predicted protein sequences of the transcripts to the embryophyta orthoDBv10 dataset (*-l embryophyta_odb10* option), consisting of 1614 conserved single copy orthologs. Protein-coding transcripts were used to supplement the computational genome annotation.

#### Methods S3: Computational genome annotation using BRAKER2

In a first run, we used as protein database 3,536,219 protein sequences from 117 Viridiplantae species from orthoDB v10 (Kriventseva *et al*. 2019), as well as the protein sequences of *Amaranthus cruentus* (Ma *et al*. 2021), as external evidence to guide the annotation. We used this preliminary run of BRAKER2 to guide mapping of the RNA-seq dataset described above to the reference genome v2.2 (Table S1). Quality control of downloaded fastq files was done using FastQC v0.11.9 (Andrews 2010). We indexed the polished reference genome using STAR v2.7.8a (Dobin and Gingeras 2015) with the following settings: *sjdbOverhang* 89, *genomeSAindexNbases* 13. We mapped the RNA-seq reads to the indexed genome using the STAR v2.7.8a (Dobin and Gingeras 2015) with default settings, using the protein-based BRAKER2 annotation as splice-junction database. To generate the final computational gene annotation, we ran BRAKER2 with reference genome v2.2, using as external evidence both the protein database as well as the previously mapped short-read RNA sequencing reads. After computational annotation of the polished genome, we extracted protein and coding sequence (CDS) fasta sequences from the BRAKER2 annotation using the AUGUSTUS script *getAnnoFastaFromJoingenes.py* (Stanke *et al*. 2006), filtering out all predicted genes with internal stop codons. We divided the predicted genes into three subcategories based on the amount of external support using the *selectSupportedSubsets.py* script of BRAKER2 (Brůna *et al*. 2021). The predicted genes were classified as “full support” if all introns of multi-exons genes were supported by external evidence (protein or RNA-seq alignments) or if both start and stop codon were supported by external evidence in the case of mono-exon genes. Genes were classified into the “any support” subset, if at least one intron or start or stop codon were supported by external evidence. Otherwise, predicted genes were classified into the “no support” subset.

#### Methods S4: Combining full-length transcripts and computational annotation

We used the supported dataset of predicted genes and the annotation file of external support hints from the BRAKER2 gene prediction together with the genomic positions of the coding sequence of the full-length transcripts, as predicted by CPC2, and the available long-read configuration file as input for TSEBRA. We used gffread v0.12.7 (Pertea and Pertea 2020) with the *-M* and *-K* options to discard redundant gene models, removing all genes which were fully contained in another gene model, to generate the genome annotation v2.2. We manually corrected the annotation of the *AmMYBl1* gene (AHp014591) based on alignment of previous complementary DNA (cDNA) Sanger sequencing results (generated from the dark seeded accession PI 604581) to the polished reference genome using minimap2 v2.17 (Li 2018). In order to estimate the completeness of the genome annotation v2.2, we used the annotated protein sequences as input for BUSCO v5.2.2 (Simão *et al*. 2015) with the Embryophyte reference set (*-l embryophyta_odb10* option). For each annotated isoform of the genome annotation v2.2, we assessed support by assembled Iso-Seq transcripts using gffcompare v0.12.6 (Pertea and Pertea 2020). To compare the annotation completeness with the previously published genome annotations of *A. hypochondriacus* and *A. cruentus*, we submitted the annotated protein sequences from the two genome annotations to BUSCO v5.2.2 using the same parameters as detailed above. We visualized the genome annotation in a circos plot using the circlize R package (Gu *et al*. 2014), calculating gene and repetitive element density in 1 Mb windows along the genome.

#### Methods S5: Functional annotation

Functional annotation of annotated protein sequences was carried out using eggNOG-mapper v2.17 (online version; Cantalapiedra *et al*. 2021, *taxonomic scope*=Viridiplantae (33090), *E-value*=0.001, *score*=60, *percentage identity*=40, *query and subject coverage*=20, transferring all non-electronic gene onthology (GO) annotations) and Mercator4 v.5.0 (Schwacke *et al*. 2019). We additionally annotated sequences using Interproscan v5.56-89.0 (Jones *et al*. 2014), retrieving pathway and GO-term annotations. We identified betalain pathway genes using BLAST (Altschul *et al*. 1997) searches with previously described betalain pathway genes (Table S3). Searches were carried out against the annotated amaranth protein sequences using BLAST+ v2.9.0-2, requiring a high-scoring query coverage of 80% (*qcov_hsp_perc 80*) and using the best hit as candidate sequence. For phylogenetic analysis of the candidate sequences, we aligned the amaranth CYP76AD and DODA protein sequences with sequences of other species (Table S9 and S10) using Clustal Omega v1.2.4 (Sievers *et al*. 2011) with default settings and constructed neighbour joining phylogenetic trees using ClustalW v2.1 (Thompson *et al*. 1994) with 1000 bootstrap replicates. We used the flavonoid pathway identification pipeline KIPEs v0.35 (Pucker *et al*. 2020) to identify candidate genes of the flavonoid pathway. Amaranth genes were defined as candidate genes if they included all of the conserved residues defined by KIPEs. If no candidate sequence with all conserved residues could be identified, instead the amaranth gene with the highest percentage similarity and less than three missing conserved residues was defined as candidate gene. Few amaranth genes represented the best candidates for multiple different flavonoid pathway genes (e.g. *F3’H* and *F3’5’H* shared the best candidate in AHp017497.1). Since these ambiguous candidate genes could be assigned to specific flavonoid pathway genes with high confidence (displaying all conserved residues), we defined the genes *CHI2*, *F3’5’H*, *FNS1* and *ANS* to be missing in amaranth.

To identify potential MYB transcription factors, we searched the predicted protein sequences of the *A. hypochondriacus* genome annotation v2.2 for matches to the MYB DNA-binding domain HMM profile (Pfam ID: PF00249, downloaded 11.01.2022) using HMMer v3.3.2 (Eddy 2009). We filtered matching domains based on score (> 25), alignment length (> 20 amino acids) and accuracy (> 0.8) and divided genes with matching domains into the R2R3, 3R and 4R subfamilies based on the number of adjacent repeats of the MYB domain using a custom script. Domains were considered adjacent, if they were located within 15 amino acids of one another. To assign the identified MYB transcription factors to their established subgroups, we aligned amaranth MYB protein sequences with the MYB proteins of *A. thaliana* (Stracke *et al*. 2001) and *B. vulgaris* (Stracke *et al*. 2014) using Clustal Omega v1.2.4 (Sievers *et al*. 2011) with default settings. We constructed a neighbour joining phylogenetic tree with 1000 bootstrap replicates from the Clustal Omega multiple sequence alignment using ClustalW v2.1 (Thompson *et al*. 1994). MYB proteins were assigned to subgroups based on the subgroup assignment of the closest *A. thaliana* homolog, as reported by Dubos *et al*. (2010) and Li *et al*. (2016). For ambiguous placement in the phylogeny, additionally the subgroup assignment of the closest *B. vulgaris* homolog was considered (Stracke *et al*. 2014). For *A. thaliana* subgroup S8, we considered the classification also used by Tombuloglu (2020). We identified potential bHLH-interacting MYB transcription factors by searches in the protein sequence using the conserved sequence motif ([D/E]Lx_2_[R/K]x_3_Lx_6_Lx_3_R), described by Zimmermann *et al*. (2004). To investigate their sequence conservation, we aligned subgroup S6 anthocyanin regulators and BvMYB1-clade proteins using ClustalOmega with default settings.

#### Methods S6: Betalain quantification

We performed photometric quantification and quantified individual metabolites (amaranthin and betanin (betacyanins), vulgaxanthin IV (betaxanthin), and betalamic acid) using liquid chromatography - mass spectrometry (LC-MS) from the same samples with at least 3 replicates per tissue per accession (4 for leaves of PI 576485, 8 for leaves of PI 538323, 3 for leaves of PI 686465, 3 for flower of PI 576485, 8 for flower of PI 538323 and 3 for flower of PI 686465). We performed additional photometric quantification from 5 leaf samples of PI 576485, 2 leaf samples of PI 538323 and 4 leaf samples of PI 686465, as well as 5 flower samples of PI 576485, 2 flower samples of PI 538323 and 5 flower samples of PI 686465. To quantify extracted betalains, we modified a previously described extraction method (Chang *et al*. 2021; von Elbe 2001). We sampled around 30 mg of flower or leaf tissue and immediately stored plant material in liquid nitrogen. Flower tissue was sampled from the tip of the inflorescense, leaf tissue from the tip of the youngest fully developed leaf. We homogenized plant material in liquid nitrogen and extracted betalains in a two step procedure. Betalains were extracted on a shaker in H_2_O for 60 min at 8 °C, afterwards samples were centrifuged at 16,000 g for 10 min at 4 °C. After centrifugation, the supernatant was transferred to a new tube and methanol and chloroform were added in a ratio of 1:1:2 H_2_O:methanol:chloroform to separate residual chlorophyll from extracted betalains. For usage as internal standard in LC-MS measurements, we added 10 µM sinigrin to methanol in the previous step. Extracts were incubated on a shaker for 10 min at 8 °C and centrifuged again at 16,000 g for 10 min at 4 °C. We collected betalains from the upper aqueous phase for quantification using photometry and LC-MS.

For photometric quantification of betacyanin and betaxanthin content, we measured absorption of the extracts at 536 nm (*A*_536_) and 480 nm (*A*_480_), respectively. We converted *A*_536_ to betacyanin content using the molar extinction coefficient *ɛ* = 60, 000 *M^−^*^1^ *cm^−^*^1^ (Jain *et al*. 2015) and the molecular weight of amaranthin (726.6 *g mol^−^*^1^). To correct for betacyanin absorption at 480 nm, we adjusted our estimation of betaxanthin absorption using the formula 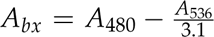 (von Elbe 2001). We converted *A_bx_* to betaxanthin content using the molar extinction coefficient *ɛ* = 48, 000 *M^−^*^1^ *cm^−^*^1^ (Escribano *et al*. 2017) and the molecular weight of vulgaxanthin IV (324.333 *g mol^−^*^1^).

LC of plant extracts was performed on the Dionex UltiMate 3000 UPLC system (Thermo Scientific, Germering, Germany) using a C18 XSelect® HSS T3 (2.5 µm, 3.0x150 mm) column (Waters, Drinagh, Ireland) at a flow rate of 0.35 ml/min. The mobile phase consisted of water/0.1 % formic acid (solution A) and methanol/0.1 % formic acid (solution B). The 35-min gradient conditions were as follows: 0-3 min 1 % B (v/v), 3-20 min linear gradient to 40 %, 20-25 min linear gradient to 99 % B, 25-30 min 99 % B, 30-33 min linear gradient to 1 % B and 33-35 min 1 % B. MS spectra data were acquired using a maXis 4G QTOF-instrument (Bruker Daltonics, Bremen, Germany) equipped with an electrospray ion source. Betalains were analyzed in positive ion mode with following source settings: capillary voltage: 3.5 kV, nebulizer pressure: 1 bar, dry gas flow: 8 l/min, dry temperature: 200 °C. Metabolite identification was achieved from full-scan MS data (mass range m/z 50-1000 m/z) using the DataAnalysis (version 6.0) software (Bruker, Bremen, Germany). Retention time of betalains was confirmed by comparisons to the literature. Peak areas of respective compounds using the extracted ion chromatograms were determined with QuantAnalysis (version 6.0) software (Bruker, Bremen, Germany). Betaxanthin and betacyanin amounts were normalized against sample fresh weight and the internal standard 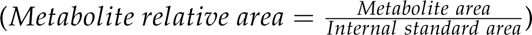. We conducted statistical analysis of betalain content differences between samples within tissue using ANOVA.

### Supplementary Results

#### Results S1: Improvement of the grain amaranth reference assembly

In order to improve the annotation with full-length transcript sequencing evidence, we performed PacBio isoform sequencing of seven different tissues of the amaranth reference accession (PI 558499, Table S1). A total of 5,306,288 CCS reads across the seven different tissues were generated, resulting in a total of 163,587 high-quality transcript sequences after filtering and clustering (Table S2). Of these high-quality transcripts, 94.23 % (154,949) could be uniquely mapped to the polished reference genome and were further collapsed into 53,638 unique isoforms. We further filtered possible 5’ degraded sequences and corrected sequences according to the polished reference genome, resulting in a set of 52,468 transcript sequences from 22,165 loci. Out of these, a total of 42,835 transcripts from 18,593 genes were predicted as coding with intact ORF. Correction of long-read transcript sequences using the polished reference genome resulted in a minor increase in reported BUSCO completeness from 90.7 % to 90.8 % (Table S8).

To evaluate the effect of genome polishing on long-read ‘error correction’, we mapped HQ long-read clustered isoforms from all tissues to the unpolished reference genome and collapsed the mapped isoforms using identical settings as above. Genome-based correction of long-read transcripts is expected to increase transcript accuracy by correcting possible sequencing errors in the long-reads with the reference genome sequence. A decreased accuracy of transcripts after genome-based correction instead indicates sequencing errors in the assembly not present in the transcripts. We ‘corrected’ the sequences of the collapsed isoforms using the unpolished reference genome and predict ORFs in both the uncorrected and corrected isoforms. For a total of 43,245 transcript sequences that were predicted to be coding before and after correction using the unpolished reference genome, 35.1 % (15,194) differed in predicted ORF length. While 12,289 transcripts had longer predicted ORFs without ‘correction’, 2,805 were predicted with longer ORF after ‘correction’ with the unpolished reference genome. ORFs of transcripts with differential ORF predictions before and after ‘correction’ differed by an average of 140 amino acids. SQANTI based correction of long-read transcripts using the unpolished reference genome strongly decreased the reported BUSCO score from 90.6 % to 74.6 % complete (Table S8). The large difference in BUSCO score and predicted ORF length between Iso-Seq data and unpolished genome indicate that the unpolished reference genome sequence included indels that disrupted the previous annotation. In contrast to that, reference genome based correction using our polished assembly resulted in an increase in reported BUSCO completeness from 90.7 % to 90.8 % (Table S8) with only 10.1 % of transcripts differing in predicted ORF length after correction. These results demonstrate the successful removal of sequencing errors from the reference genome through genome polishing and the positive impact for gene annotation.

#### Results S2: Annotation of the polished A. hypochondriacus reference genome increases completeness of annotated genes

In addition to the long-read RNA annotation, we predicted protein-coding genes in the polished reference genome using BRAKER2 (Brůna *et al*. 2021) with external evidence in the form of a protein database and short-read RNA sequencing reads from eight different tissues (Table S1). Computational gene prediction resulted in a total of 32,604 predicted gene isoforms, 67.5 % (22,012) with full support, 21.5 % (7,013) with partial support and 11 % (3,579) without external support. We excluded all predicted isoforms without support from RNA sequencing or protein alignments. The filtered computational annotation included 29,025 annotated isoforms from 25,405 genes with a BUSCO score of 97.9 % complete (Table S8).

After combination of BRAKER2 and Iso-Seq annotations, we found the complete intron chain of 20,959 (74.66 %) annotated isoforms to be supported by full-length transcripts from long-read sequencing data, indicating high-quality of the annotation. As expected, regions with high repeat content show less annotated genes (Fig. 1c). While the number of predicted genes was comparable to the previously published annotation (Lightfoot *et al*. 2017) and lower than the number of predicted genes from *A. cruentus* (Ma *et al*. 2021), the mean CDS length and exon number was slightly increased (Table 1). While the previous *A. hypochondriacus* genome annotation included only a single primary isoform for each gene, our genome annotation v2.2 includes multiple annotated isoforms for 3,484 genes, resulting in a total of 28,074 annotated isoforms.

### Supplementary Tables

**Table S1.** Summary of tissue of origin, read count, and source of long-read and short-read RNA sequencing data used to generate genome annotation v2.2. For long-read data, the number of reads refers to CCS reads. For short-read data, the number of reads refers to read pairs.

| Tissue | Sequencing type | Reads | Uniquely mapped (%) | Source |
| --- | --- | --- | --- | --- |
| Pollen | long-read | 637,760 | - | this study |
| Root | long-read | 752,123 | - | this study |
| Root | short-read | 22,656,771 | 93.13 | Clouse *et al.* (2016) |
| Cotyledons | long-read | 928,752 | - | this study |
| Cotyledons | short-read | 22,291,775 | 89.50 | Clouse *et al.* (2016) |
| Flower | long-read | 691,694 | - | this study |
| Flower | short-read | 21,501,431 | 94.93 | Clouse *et al.* (2016) |
| Leaf | long-read | 478,352 | - | this study |
| Leaf | short-read | 22,045,070 | 94.28 | Clouse *et al.* (2016) |
| Developing seed | long-read | 371,693 | - | this study |
| Developing seed | short-read | 21,768,342 | 96.31 | Clouse *et al.* (2016) |
| Mature seed | long-read | 1,445,914 | - | this study |
| Mature seed | short-read | 22,242,512 | 92.17 | Clouse *et al.* (2016) |
| Stem | short-read | 22,162,620 | 95.61 | Clouse *et al.* (2016) |
| Water-stressed tissues | short-read | 21,610,476 | 95.37 | Clouse *et al.* (2016) |

**Table S2.** Number of PacBio Iso-Seq CCS and FLNC reads for all sequenced tissues. We combined the FLNC reads from all tissues before clustering FLNC reads into a set of HQ isoforms from all tissues.

|  | CCS reads | FLNC reads | HQ clustered |
| --- | --- | --- | --- |
| Root | 752,123 | 734,212 |  |
| Cotyledon | 928,752 | 895,195 |  |
| Flower | 691,694 | 676,684 |  |
| Leaf | 478,352 | 467,382 |  |
| Pollen | 637,760 | 624,434 |  |
| Developing seed | 371,693 | 363,574 |  |
| Mature seed | 1,445,914 | 1,404,972 |  |
| Combined | 5,306,288 | 5,166,453 | 163,587 |

**Table S3.** Summary of BLAST results for betalain pathway genes. GenBank or Uniprot entries for query genes are noted alongside their original publication. For each query, the best hit in the annotated *A. hypochondriacus* protein sequences and the percentage identity as reported by blast is displayed.

| Pathway gene | Query GenBank | Query Uniprot | Reference | Query species | Best hit | Percentage identity |
| --- | --- | --- | --- | --- | --- | --- |
| CYP76AD1 | - | C76AD_BETVU | Hatlestad et al. (2012) | *B. vulgaris* | AHp023148 | 84.14 |
| CYP76AD2 | HQ656025.1 | - | Hatlestad et al. (2012) | *A. cruentus* | AHp023148 | 99.80 |
| CYP76AD5 | - | A0A0U1YLX5_BETVU | Sunnadeniya et al. (2016) | *B. vulgaris* | AHp000674 | 83.43 |
| CYP76AD6 | - | A0A0U1YJA0_BETVU | Sunnadeniya et al. (2016) | *B. vulgaris* | AHp000674 | 82.16 |
| DODA1 | HQ889614.1 | - | Casique-Arroyo et al. (2014) | *A. hypochondriacus* | AHp010386 | 100.00 |
| DODA1 | - | DOD1U_BETVU | Hatlestad et al. (2012) | *B. vulgaris* | AHp023147 | 78.10 |
| cDOPA5GT | KJ136018.1 | - | Casique-Arroyo et al. (2014) | *A. hypochondriacus* | AHp007219 | 100.00 |
| Betanidin5GT | KJ136017.1 | - | Casique-Arroyo et al. (2014) | *A. hypochondriacus* | AHp001663 | 99.79 |
| Betanidin6GT | KP174812.1 | - | Zheng et al. (2016) | *A. tricolor* | AHp005940 | 92.96 |

**Table S4.** Statistics for whole-genome sequencing data used in the BSAs. The number of read pairs generated for the green and red bulks of both BSAs and the percentage of read pairs uniquely mapped to the polished reference genome is displayed.

| Sample | Reads | Uniquely mapped (%) |
| --- | --- | --- |
| green bulk regulator BSA | 59,471,388 | 98.29 |
| red bulk regulator BSA | 68,365,577 | 98.62 |
| green bulk biosynthesis BSA | 63,069,713 | 98.03 |
| red bulk biosynthesis BSA | 71,253,748 | 97.99 |

**Table S5.** Statistics for RNA sequencing data generated from BSA bulks. Number of read pairs before and after adapter trimming for short-read RNA-seq data generated for gene expression quantification in pooled green and red flower tissue of regulator and biosynthesis BSA. The data was generated from pooled flower tissue of 24 plants from each bulk of the biosynthesis BSA and 29 plants from each bulk of the regulator BSA.

| Sample | Reads | Trimmed reads |
| --- | --- | --- |
| green bulk regulator BSA | 25,960,085 | 25,940,297 |
| red bulk regulator BSA | 25,423,535 | 25,404,128 |
| green bulk biosynthesis BSA | 38,659,017 | 38,560,213 |
| red bulk biosynthesis BSA | 25,718,709 | 25,673,931 |

**Table S6.**
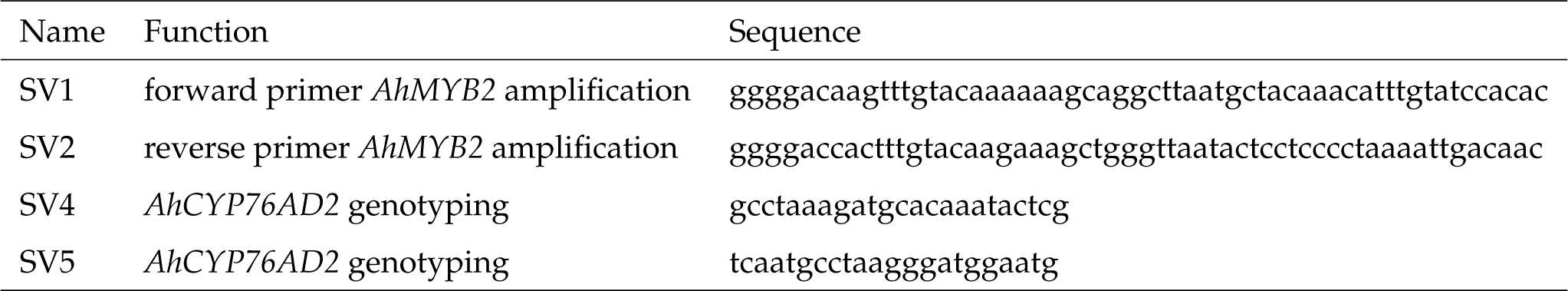
List of primers used in the study.

**Table S7.** Repetitive element content of the polished *A. hypochondriacus* genome assembly v2.2.

| Classification | Genome covered [bp] | Genome covered [%] |
| --- | --- | --- |
| DNA/CMC-EnSpm | 4,098,198 | 1.01 |
| DNA/MULE-MuDR | 5,723,062 | 1.42 |
| DNA/PIF-Harbinger | 1,168,228 | 0.29 |
| DNA/TcMar-Stowaway | 1,239,206 | 0.31 |
| DNA/Zisupton | 800,423 | 0.2 |
| DNA/hAT-Ac | 2,943,534 | 0.73 |
| DNA/hAT-Tag1 | 254,129 | 0.06 |
| DNA/hAT-Tip100 | 1,342,775 | 0.33 |
| RC/Helitron | 2,071,001 | 0.51 |
| LINE/CRE | 1,014,983 | 0.25 |
| LINE/L1 | 8,821,523 | 2.18 |
| LINE/Penelope | 21,568 | 0.01 |
| LINE/R1 | 18,480 | 0 |
| LINE/RTE | 504,828 | 0.12 |
| LINE/RTE-BovB | 2,580,488 | 0.64 |
| LTR | 163,433 | 0.04 |
| LTR/Copia | 39,486,653 | 9.77 |
| LTR/Gypsy | 19,135,979 | 4.74 |
| LTR/Pao | 50,089 | 0.01 |
| LTR/Unknown | 10,497,276 | 2.6 |
| SINE/5S | 76,086 | 0.02 |
| SINE/U | 39,697 | 0.01 |
| SINE/tRNA | 697,466 | 0.17 |
| SINE/tRNA-CR1 | 38,625 | 0.01 |
| rRNA | 21,280 | 0.01 |
| tRNA | 30,256 | 0.01 |
| Low complexity | 937,694 | 0.23 |
| Simple repeat | 11,495,793 | 2.85 |
| Unknown | 114,642,357 | 28.38 |
| Total sum | 229,915,110 | 56.91 |

**Table S8.** BUSCO score summary of different data sets against the Embryophyta reference set. Iso-Seq FLNC reads were mapped against the unpolished or polished reference genome and corrected using the unpolished or polished reference genome sequence. For Iso-Seq transcript sets, BUSCO was run in protein mode using predicted ORFs. For genome annotations, BUSCO was run in protein mode on predicted protein sequences.

| Dataset | Species | Data source | Complete | Complete (single copy) | Complete (duplicated) | Fragmented | Missing |
| --- | --- | --- | --- | --- | --- | --- | --- |
| Genome annotation v2.1 | *A. hypochondriacus* | Lightfoot *et al.* (2017) | 80.7 | 78.5 | 2.2 | 8.8 | 10.5 |
| Genome annotation | *A. cruentus* | Ma *et al.* (2021) | 89.8 | 87.2 | 2.6 | 2.5 | 7.7 |
| Iso-Seq transcripts (uncorrected) | *A. hypochondriacus* | this study | 90.7 | 58.9 | 31.8 | 2.0 | 7.3 |
| Iso-Seq transcripts (corrected) | *A. hypochondriacus* | this study | 90.8 | 57.3 | 33.5 | 1.4 | 7.8 |
| Iso-Seq transcripts (unpolished genome, uncorrected) | *A. hypochondriacus* | this study | 90.6 | 58.7 | 31.9 | 2.0 | 7.4 |
| Iso-Seq transcripts (unpolished genome, corrected) | *A. hypochondriacus* | this study | 74.6 | 46.7 | 27.9 | 11.0 | 14.4 |
| Braker2 annotation | *A. hypochondriacus* | this study | 97.9 | 80.9 | 17.0 | 0.8 | 1.3 |
| Genome annotation v2.2 | *A. hypochondriacus* | this study | 98.0 | 83.4 | 14.6 | 0.7 | 1.3 |

**Table S9.** Sequences used for phylogenetic analysis of amaranth CYP76AD genes. Species, gene name and GenBank accession of input genes for phylogenetic analysis.

| Species | Gene name | GenBank accession |
| --- | --- | --- |
| *Arabidopsis thaliana* | AtCYP76C2 | NP_182081 |
| *Arabidopsis thaliana* | AtCYP94B3 | NP_190421 |
| *Basella alba* | BaCYP76AD13 | AJD87469 |
| *Bougainvillea peruviana* | BpCYP76AD1-1 | BCD59210 |
| *Bougainvillea peruviana* | BpCYP76AD1-2 | BCD59211 |
| *Bougainvillea peruviana* | BpCYP76AD1-3 | BCD59212 |
| *Bougainvillea peruviana* | BpCYP76AD2 | BCD59213 |
| *Beta vulgaris* | BvCYP76AD1 | AET43290 |
| *Beta vulgaris* | BvCYP76AD5 | AJD87473 |
| *Beta vulgaris* | BvCYP76AD6 | AJD87474 |
| *Chenopodium quinoa* | CqCYP76AD1 | XP_021769302 |
| *Mirabilis jalapa* | MjCYP76AD3 | AET43292 |
| *Mirabilis jalapa* | MjCYP76AD7 | AJD87463 |
| *Mirabilis jalapa* | MjCYP76AD15 | AJD87471 |

**Table S10.** Sequences used for phylogenetic analysis of amaranth DODA genes. Species, gene name and GenBank accession of input genes for phylogenetic analysis.

| Species | Gene name | GenBank accession |
| --- | --- | --- |
| Beta vulgaris | BvDODAα1 | MN153194 |
| Beta vulgaris | BvDODAα2 | MN153195 |
| Beta vulgaris | BvDODAβ | MN153198 |
| Chenopodium quinoa | CqDODA1α1 | XP_021769303 |
| Mirabilis jalapa | MjDOD | AB435372.1 |
| Mesembryanthemum crystallinum | McDODAα 1 | MN136225 |
| Mesembryanthemum crystallinum | McDODAβ | MN136227 |

**Table S11.** Amaranth flavonoid pathway candidate genes identified using KIPEs. Displayed are candidate gene names from genome annotation v2.2 alongside the percentage similarity, the percentage of conserved residues and the number of missing conserved residues as reported by KIPEs. Missing pathway genes (defined by having no unique candidate genes or three or more missing conserved residues) were marked with asterisks and defined as unidentified in *A. hypochondriacus*.

| Pathway gene | Candidate gene | Percentage similarity | Percentage conserved residues | Missing conserved residues |
| --- | --- | --- | --- | --- |
| PAL | AHp012752 | 79.49 | 100.0 | 0 |
| PAL | AHp021980 | 79.40 | 100.0 | 0 |
| C4H | AHp013217 | 81.06 | 100.0 | 0 |
| C4H | AHp022384 | 80.66 | 100.0 | 0 |
| C4H | AHp022382 | 72.71 | 100.0 | 0 |
| 4CL | AHp014409 | 64.73 | 100.0 | 0 |
| 4CL | AHp020962 | 65.36 | 100.0 | 0 |
| CHS | AHp004305 | 79.78 | 94.29 | 2 |
| CHI1 | AHp009962 | 59.97 | 100.0 | 0 |
| CHI2* | no candidate | - | - | - |
| F3'H | AHp017497 | 68.15 | 100.0 | 0 |
| F3'H | AHp022122 | 36.88 | 100.0 | 0 |
| F3'H | AHp003152 | 38.80 | 100.0 | 0 |
| F3'H | AHp022120 | 37.24 | 100.0 | 0 |
| F3'H | AHp022123 | 25.78 | 100.0 | 0 |
| F3'5'H* | no candidate | - | - | - |
| FNS1* | no candidate | - | - | - |
| DFR | AHp009303 | 61.99 | 96.43 | 1 |
| F3H | AHp006454 | 79.68 | 100.0 | 0 |
| FLS | AHp008991 | 62.03 | 100.0 | 0 |
| LAR | AHp017409 | 57.70 | 100.0 | 0 |
| ANS* | no candidate | - | - | - |
| ANR* | AHp001635 | 37.14 | 86.96 | 3 |

**Table S12.** MYB transcription factors identified in *A. hypochondriacus*. Genes were assigned into subgroups based on their phylogenetic placement in comparison to *A. thaliana* and *B. vulgaris* MYBs. Subgroup functions were assigned according to conserved subgroup functions (see Materials and Methods).

| Gene name | Gene id | Transcript id | Scaffold | Start | End | Strand | Subfamily | Protein length | Exon number | Subgroup | Functional assignment |
| --- | --- | --- | --- | --- | --- | --- | --- | --- | --- | --- | --- |
| - | AHp000437 | AHp000437.1 | Scaffold_1 | 4347523 | 4351157 | + | R2R3 | 270 | 3 | S16 | Photomorphogenesis |
| - | AHp000789 | AHp000789.1 | Scaffold_1 | 10090637 | 10092364 | + | R2R3 | 255 | 2 | S4 | Repressors of phenylpropanoids, sinapate, lignin |
| - | AHp001699 | AHp001699.1 | Scaffold_1 | 29115865 | 29116587 | + | R2R3 | 241 | 1 | S22 | Defense, stress response |
| - | AHp001699 | AHp001699.2 | Scaffold_1 | 29115865 | 29116656 | + | R2R3 | 239 | 2 | S22 | Defense, stress response |
| - | AHp002277 | AHp002277.1 | Scaffold_1 | 35637528 | 35639576 | - | R2R3 | 272 | 2 | S4 | Repressors of phenylpropanoids, sinapate, lignin |
| - | AHp002664 | AHp002664.1 | Scaffold_2 | 1693532 | 1697261 | + | R2R3 | 274 | 3 | S2 | Abiotic stress response, flower morphogenesis |
| - | AHp002686 | AHp002686.1 | Scaffold_2 | 1885905 | 1892082 | - | R2R3 | 317 | 3 | S7 | Flavonol, flavonoid pathway |
| - | AHp002839 | AHp002839.1 | Scaffold_2 | 3368079 | 3370731 | + | R2R3 | 322 | 3 | S1 | Defense, stress response |
| - | AHp003887 | AHp003887.1 | Scaffold_2 | 26854106 | 26855787 | + | R2R3 | 309 | 3 | S14 | Axillary meristem, root growth |
| - | AHp003952 | AHp003952.1 | Scaffold_2 | 27861374 | 27862381 | - | R2R3 | 336 | 1 | S22 | Defense, stress response |
| - | AHp004623 | AHp004623.1 | Scaffold_2 | 34693851 | 34698394 | - | R2R3 | 454 | 12 | - | - |
| - | AHp005058 | AHp005058.1 | Scaffold_3 | 3408139 | 3411828 | + | R2R3 | 359 | 4 | S21 | Cell wall, lignin, axillary meristem |
| - | AHp005110 | AHp005110.1 | Scaffold_3 | 3994528 | 4000152 | + | R2R3 | 454 | 12 | - | - |
| - | AHp005488 | AHp005488.1 | Scaffold_3 | 8254983 | 8258030 | + | R2R3 | 269 | 3 | S20 | Defense, stress response |
| - | AHp005948 | AHp005948.1 | Scaffold_3 | 14823654 | 14827853 | - | R2R3 | 307 | 3 | - | - |
| - | AHp006062 | AHp006062.1 | Scaffold_3 | 18197344 | 18199249 | - | R2R3 | 171 | 4 | S24 | - |
| - | AHp006213 | AHp006213.1 | Scaffold_3 | 24208195 | 24209124 | + | R2R3 | 310 | 1 | S22 | Defense, stress response |
| - | AHp006562 | AHp006562.1 | Scaffold_4 | 605153 | 607206 | + | R2R3 | 316 | 3 | S1 | Defense, stress response |
| - | AHp006562 | AHp006562.2 | Scaffold_4 | 605153 | 608233 | + | R2R3 | 178 | 4 | S1 | Defense, stress response |
| - | AHp007248 | AHp007248.1 | Scaffold_4 | 11044442 | 11046030 | + | R2R3 | 359 | 3 | S10 | - |
| - | AHp007516 | AHp007516.1 | Scaffold_4 | 18919997 | 18922556 | + | R2R3 | 540 | 3 | S18 | Anther development, stress response |
| - | AHp007516 | AHp007516.2 | Scaffold_4 | 18919997 | 18922556 | + | R2R3 | 535 | 3 | S18 | Anther development, stress response |
| - | AHp007908 | AHp007908.1 | Scaffold_4 | 24322377 | 24324732 | - | R2R3 | 296 | 3 | S4 | Repressors of phenylpropanoids, sinapate, lignin |
| - | AHp008229 | AHp008229.1 | Scaffold_4 | 27535327 | 27540827 | - | R2R3 | 353 | 3 | S18 | Anther development, stress response |
| - | AHp008229 | AHp008229.2 | Scaffold_4 | 27535327 | 27540827 | - | R2R3 | 375 | 4 | S18 | Anther development, stress response |
| - | AHp008229 | AHp008229.3 | Scaffold_4 | 27536451 | 27540827 | - | R2R3 | 310 | 2 | S18 | Anther development, stress response |
| - | AHp008362 | AHp008362.1 | Scaffold_5 | 579584 | 581724 | + | R2R3 | 223 | 4 | S14 | Axillary meristem, root growth |
| - | AHp008362 | AHp008362.2 | Scaffold_5 | 579584 | 581724 | + | R2R3 | 345 | 3 | S14 | Axillary meristem, root growth |
| - | AHp008410 | AHp008410.1 | Scaffold_5 | 1052626 | 1053435 | - | R2R3 | 270 | 1 | S22 | Defense, stress response |
| - | AHp008675 | AHp008675.1 | Scaffold_5 | 4050096 | 4052532 | + | R2R3 | 348 | 2 | S21 | Cell wall, lignin, axillary meristem |
| - | AHp009283 | AHp009283.1 | Scaffold_5 | 19207042 | 19209867 | + | R2R3 | 301 | 3 | S9 | Trichome branching, petal morphogenesis |
| - | AHp009452 | AHp009452.1 | Scaffold_5 | 21916376 | 21917921 | + | R2R3 | 315 | 3 | S4 | Repressors of phenylpropanoids, sinapate, lignin |
| - | AHp009484 | AHp009484.1 | Scaffold_5 | 22269940 | 22274870 | + | R2R3 | 305 | 3 | S3 | Phenylpropanoid pathway, lignin |
| - | AHp009565 | AHp009565.1 | Scaffold_5 | 23545578 | 23547934 | + | R2R3 | 348 | 2 | AtMYB5-like | Development, proanthocyanidin biosynthesis |
| - | AHp009926 | AHp009926.1 | Scaffold_6 | 2944929 | 2948000 | + | R2R3 | 330 | 3 | - | - |
| - | AHp010652 | AHp010652.1 | Scaffold_6 | 17732890 | 17739448 | - | R2R3 | 974 | 4 | - | - |
| - | AHp010658 | AHp010658.1 | Scaffold_6 | 17860532 | 17862437 | + | R2R3 | 420 | 3 | S13 | Mucilage, lignin, stomatal closure |
| - | AHp011052 | AHp011052.1 | Scaffold_6 | 22330077 | 22334786 | - | R2R3 | 411 | 2 | S16 | Photomorphogenesis |
| - | AHp011153 | AHp011153.1 | Scaffold_6 | 23290312 | 23291464 | - | R2R3 | 300 | 2 | S18 | Anther development, stress response |
| - | AHp011578 | AHp011578.1 | Scaffold_7 | 10152532 | 10155066 | - | R2R3 | 334 | 3 | - | - |
| - | AHp011578 | AHp011578.2 | Scaffold_7 | 10153374 | 10155066 | - | R2R3 | 334 | 2 | - | - |
| - | AHp011866 | AHp011866.1 | Scaffold_7 | 16613289 | 16616126 | + | R2R3 | 314 | 3 | S20 | Defense, stress response |
| - | AHp012381 | AHp012381.1 | Scaffold_7 | 23842541 | 23847434 | + | R2R3 | 254 | 3 | S16 | Photomorphogenesis |
| - | AHp012382 | AHp012382.1 | Scaffold_7 | 23851791 | 23855945 | - | R2R3 | 289 | 3 | S16 | Photomorphogenesis |
| - | AHp012383 | AHp012383.1 | Scaffold_7 | 23881163 | 23885856 | + | R2R3 | 250 | 3 | S9 | Trichome branching, petal morphogenesis |
| - | AHp012732 | AHp012732.1 | Scaffold_8 | 3120126 | 3122087 | + | R2R3 | 375 | 3 | S14 | Axillary meristem, root growth |
| - | AHp013131 | AHp013131.1 | Scaffold_8 | 6814523 | 6815510 | + | R2R3 | 255 | 3 | S17 | - |
| - | AHp013326 | AHp013326.1 | Scaffold_8 | 9237650 | 9242324 | - | R2R3 | 403 | 3 | - | - |
| - | AHp013775 | AHp013775.1 | Scaffold_8 | 16914029 | 16918169 | - | R2R3 | 411 | 3 | S25 | Embryogenesis, seed maturation |
| - | AHp014406 | AHp014406.1 | Scaffold_9 | 12055158 | 12058160 | + | R2R3 | 350 | 3 | S14 | Axillary meristem, root growth |
| - | AHp014458 | AHp014458.1 | Scaffold_9 | 12904501 | 12907181 | - | R2R3 | 242 | 3 | S14 | Axillary meristem, root growth |
| AmMYB1 | AHp014591 | AHp014591.1 | Scaffold_9 | 14711535 | 14714080 | + | R2R3 | 283 | 4 | S5 | Proanthocyanidin biosynthesis |
| - | AHp014665 | AHp014665.1 | Scaffold_9 | 15476010 | 15477107 | - | R2R3 | 366 | 1 | - | - |
| - | AHp015175 | AHp015175.1 | Scaffold_9 | 20978046 | 20979995 | - | R2R3 | 392 | 3 | S11 | Defense, stress response |
| - | AHp015176 | AHp015176.1 | Scaffold_9 | 20993200 | 20994999 | - | R2R3 | 359 | 3 | S11 | Defense, stress response |
| - | AHp015766 | AHp015766.1 | Scaffold_10 | 12455733 | 12458734 | + | R2R3 | 351 | 3 | S1 | Defense, stress response |
| - | AHp015766 | AHp015766.2 | Scaffold_10 | 12455733 | 12458734 | + | R2R3 | 341 | 3 | S1 | Defense, stress response |
| - | AHp015783 | AHp015783.1 | Scaffold_10 | 12825187 | 12827217 | - | R2R3 | 452 | 3 | S9 | Trichome branching, petal morphogenesis |
| - | AHp016111 | AHp016111.1 | Scaffold_10 | 16835722 | 16838292 | - | R2R3 | 325 | 3 | S14 | Axillary meristem, root growth |
| - | AHp016387 | AHp016387.1 | Scaffold_10 | 20282177 | 20286204 | + | R2R3 | 406 | 2 | S23 | - |
| - | AHp016456 | AHp016456.1 | Scaffold_10 | 21051032 | 21053433 | + | R2R3 | 318 | 4 | - | - |
| AhMYB3 | AHp016530 | AHp016530.1 | Scaffold_10 | 21762621 | 21764924 | - | R2R3 | 216 | 3 | BvMYB1-like | Betalain biosynthesis |
| AhMYB4 | AHp016531 | AHp016531.1 | Scaffold_10 | 21780565 | 21783050 | - | R2R3 | 228 | 3 | BvMYB1-like | Betalain biosynthesis |
| - | AHp016620 | AHp016620.1 | Scaffold_10 | 22596330 | 22599342 | - | R2R3 | 341 | 2 | S16 | Photomorphogenesis |
| - | AHp017558 | AHp017558.1 | Scaffold_11 | 17970779 | 17972903 | + | R2R3 | 293 | 3 | S1 | Defense, stress response |
| - | AHp017657 | AHp017657.1 | Scaffold_11 | 18887019 | 18889370 | - | R2R3 | 289 | 3 | S20 | Defense, stress response |
| - | AHp017658 | AHp017658.1 | Scaffold_11 | 18908562 | 18910154 | - | R2R3 | 248 | 3 | S20 | Defense, stress response |
| - | AHp017689 | AHp017689.1 | Scaffold_11 | 19220858 | 19226127 | - | R2R3 | 341 | 3 | S7 | Flavonol, flavonoid pathway |
| - | AHp017915 | AHp017915.1 | Scaffold_11 | 21654178 | 21657013 | - | R2R3 | 362 | 3 | - | - |
| - | AHp017946 | AHp017946.1 | Scaffold_11 | 22019498 | 22021190 | - | R2R3 | 294 | 3 | S25 | Embryogenesis, seed maturation |
| - | AHp017979 | AHp017979.1 | Scaffold_12 | 139570 | 144579 | - | R2R3 | 358 | 3 | - | - |
| - | AHp018270 | AHp018270.1 | Scaffold_12 | 3175247 | 3177848 | - | R2R3 | 188 | 3 | S5 | Proanthocyanidin biosynthesis |
| - | AHp018383 | AHp018383.1 | Scaffold_12 | 4698125 | 4699671 | - | R2R3 | 346 | 2 | - | - |
| - | AHp018492 | AHp018492.1 | Scaffold_12 | 5906028 | 5907870 | - | R2R3 | 306 | 2 | S8 | Lignin |
| - | AHp018627 | AHp018627.1 | Scaffold_12 | 7669283 | 7672257 | + | 3R | 513 | 7 | 3R | - |
| - | AHp019036 | AHp019036.1 | Scaffold_12 | 15482171 | 15486341 | - | R2R3 | 356 | 3 | S21 | Cell wall, lignin, axillary meristem |
| - | AHp019040 | AHp019040.1 | Scaffold_12 | 15614348 | 15629179 | + | 4R | 900 | 13 | 4R | - |
| - | AHp019040 | AHp019040.2 | Scaffold_12 | 15614348 | 15629179 | + | 4R | 902 | 13 | 4R | - |
| - | AHp019088 | AHp019088.1 | Scaffold_12 | 16532558 | 16544063 | + | 3R | 541 | 7 | 3R | - |
| - | AHp019259 | AHp019259.1 | Scaffold_12 | 20323254 | 20324899 | + | R2R3 | 313 | 2 | - | - |
| - | AHp019341 | AHp019341.1 | Scaffold_12 | 21674277 | 21675477 | - | R2R3 | 335 | 3 | - | - |
| - | AHp019341 | AHp019341.2 | Scaffold_12 | 21674277 | 21675477 | - | R2R3 | 325 | 3 | - | - |
| - | AHp020198 | AHp020198.1 | Scaffold_13 | 17051563 | 17053916 | + | R2R3 | 259 | 3 | S17 | - |
| - | AHp020524 | AHp020524.1 | Scaffold_13 | 20104457 | 20106612 | - | R2R3 | 284 | 3 | S1 | Defense, stress response |
| - | AHp020968 | AHp020968.1 | Scaffold_14 | 12366084 | 12368376 | - | R2R3 | 285 | 3 | S14 | Axillary meristem, root growth |
| - | AHp021016 | AHp021016.1 | Scaffold_14 | 13270874 | 13277249 | + | R2R3 | 412 | 3 | S25 | Embryogenesis, seed maturation |
| - | AHp021152 | AHp021152.1 | Scaffold_14 | 14968799 | 14969911 | - | R2R3 | 371 | 1 | - | - |
| - | AHp021566 | AHp021566.1 | Scaffold_14 | 19210847 | 19212496 | - | R2R3 | 381 | 3 | S11 | Defense, stress response |
| - | AHp021567 | AHp021567.1 | Scaffold_14 | 19222097 | 19225148 | - | R2R3 | 381 | 3 | S11 | Defense, stress response |
| - | AHp021873 | AHp021873.1 | Scaffold_15 | 1897249 | 1899908 | - | R2R3 | 278 | 3 | S17 | - |
| - | AHp022482 | AHp022482.1 | Scaffold_15 | 10193355 | 10194334 | + | R2R3 | 260 | 3 | S17 | - |
| - | AHp022746 | AHp022746.1 | Scaffold_16 | 532237 | 533511 | - | R2R3 | 274 | 3 | - | - |
| AhMYB2 | AHp022773 | AHp022773.1 | Scaffold_16 | 988734 | 991447 | + | R2R3 | 236 | 3 | BvMYB1-like | Betalain biosynthesis |
| AhMYB2 | AHp022773 | AHp022773.2 | Scaffold_16 | 988734 | 991447 | + | R2R3 | 240 | 3 | BvMYB1-like | Betalain biosynthesis |
| - | AHp022995 | AHp022995.1 | Scaffold_16 | 3273279 | 3276051 | - | R2R3 | 422 | 2 | S23 | - |
| - | AHp023085 | AHp023085.1 | Scaffold_16 | 4378346 | 4379815 | + | R2R3 | 243 | 2 | - | - |
| - | AHp023286 | AHp023286.1 | Scaffold_16 | 7455684 | 7465379 | - | 3R | 1017 | 11 | 3R | - |

### Supplementary Figures

**Figure S1.**
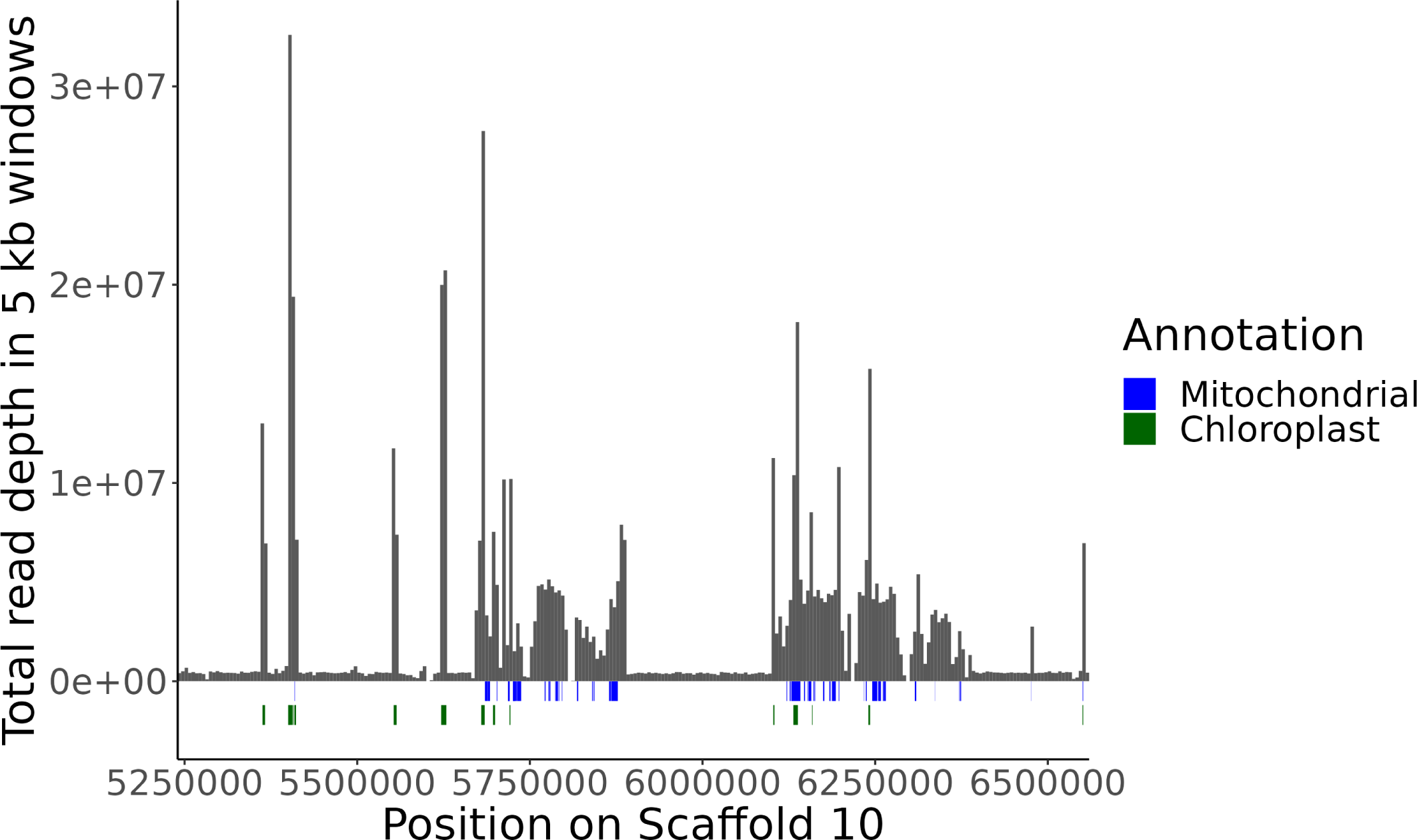
Organellar contamination in the *A. hypochondriacus* assembly. Total read depth of whole genome sequencing data in 5 kb windows along Scaffold 10 region with mapping bias. High read depth regions correspond partially to mapped chloroplast and mitochondrial regions in the genome.

**Figure S2.**
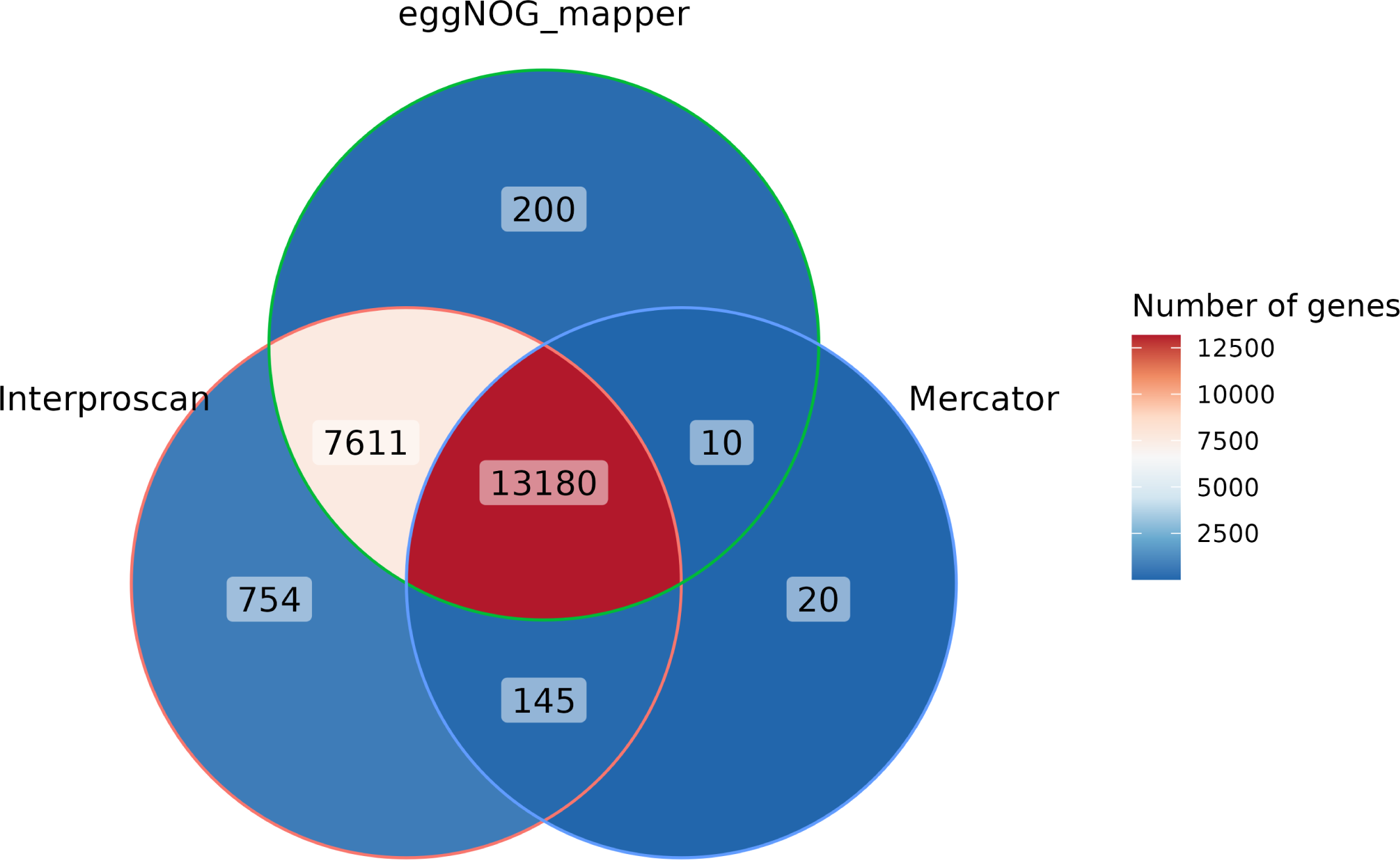
Summary of the functional annotation of the reference genome. Overlap between functionally annotated genes using the annotation tools Interproscan (considering only genes with identified Pfam, PANTHER or CDD domains as annotated), eggNOG-mapper and Mercator.

**Figure S3.**
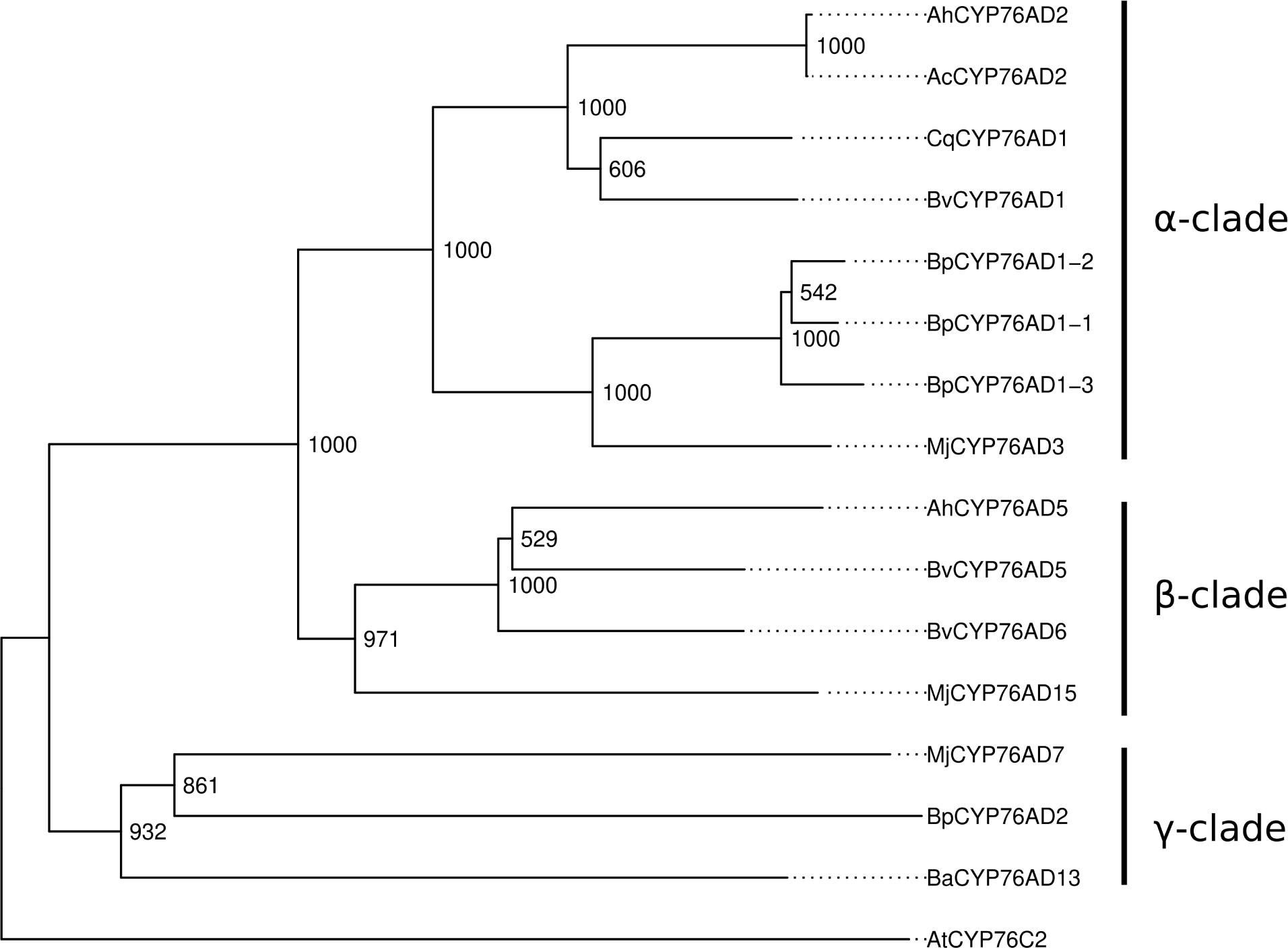
Phylogenetic analysis of amaranth CYP76AD proteins with CYP protein sequences from other species. CYP76AD *α*-, *β*- and *γ*-clades are annotated in the alignment. Bootstrap values from 1000 replicates are depicted on the tree.

**Figure S4.**
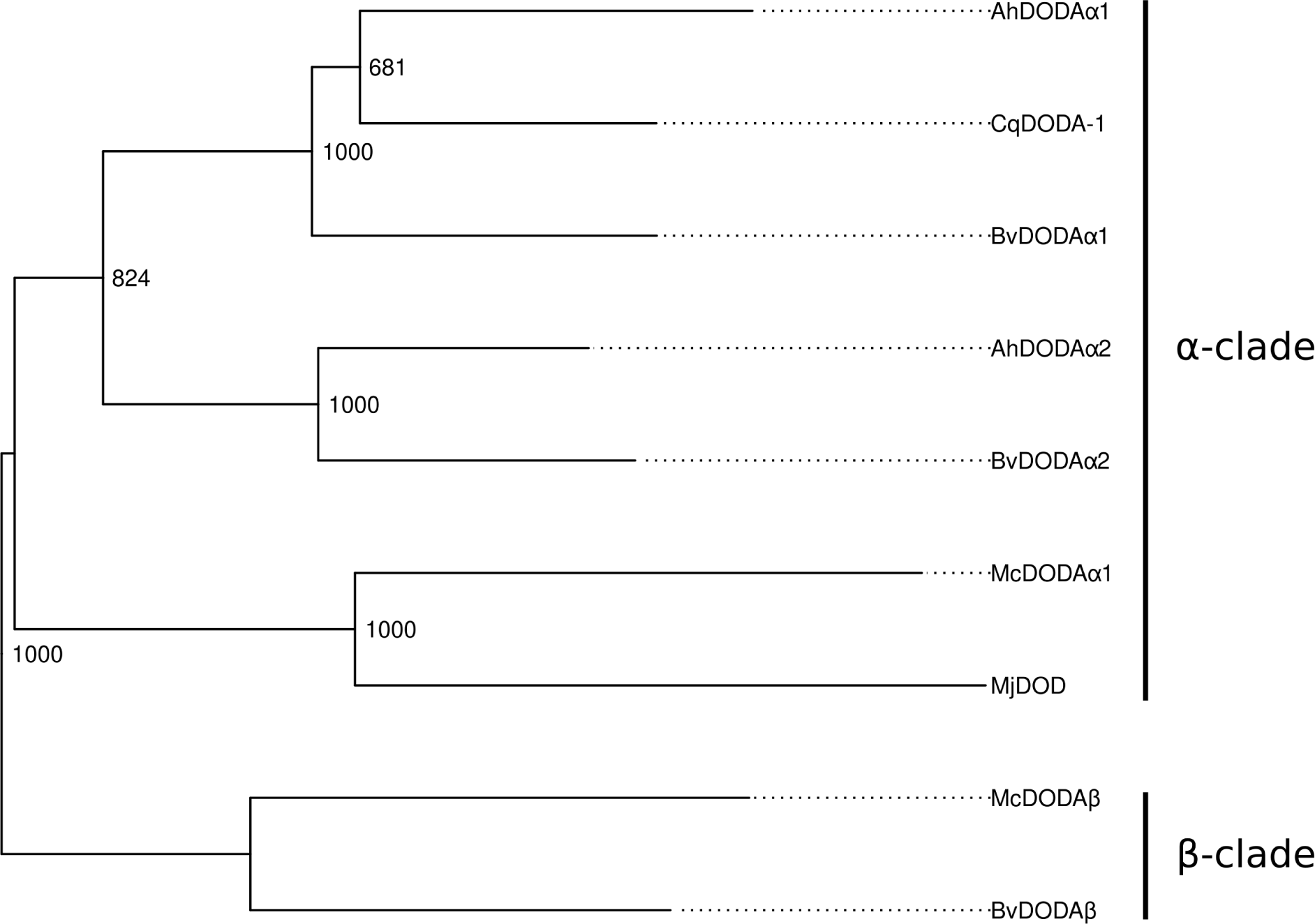
Phylogenetic analysis of amaranth DODA proteins with DODA protein sequences from other species. DODA *α*- and *β*-clades are annotated in the alignment. Bootstrap values from 1000 replicates are depicted on the tree.

**Figure S5.**
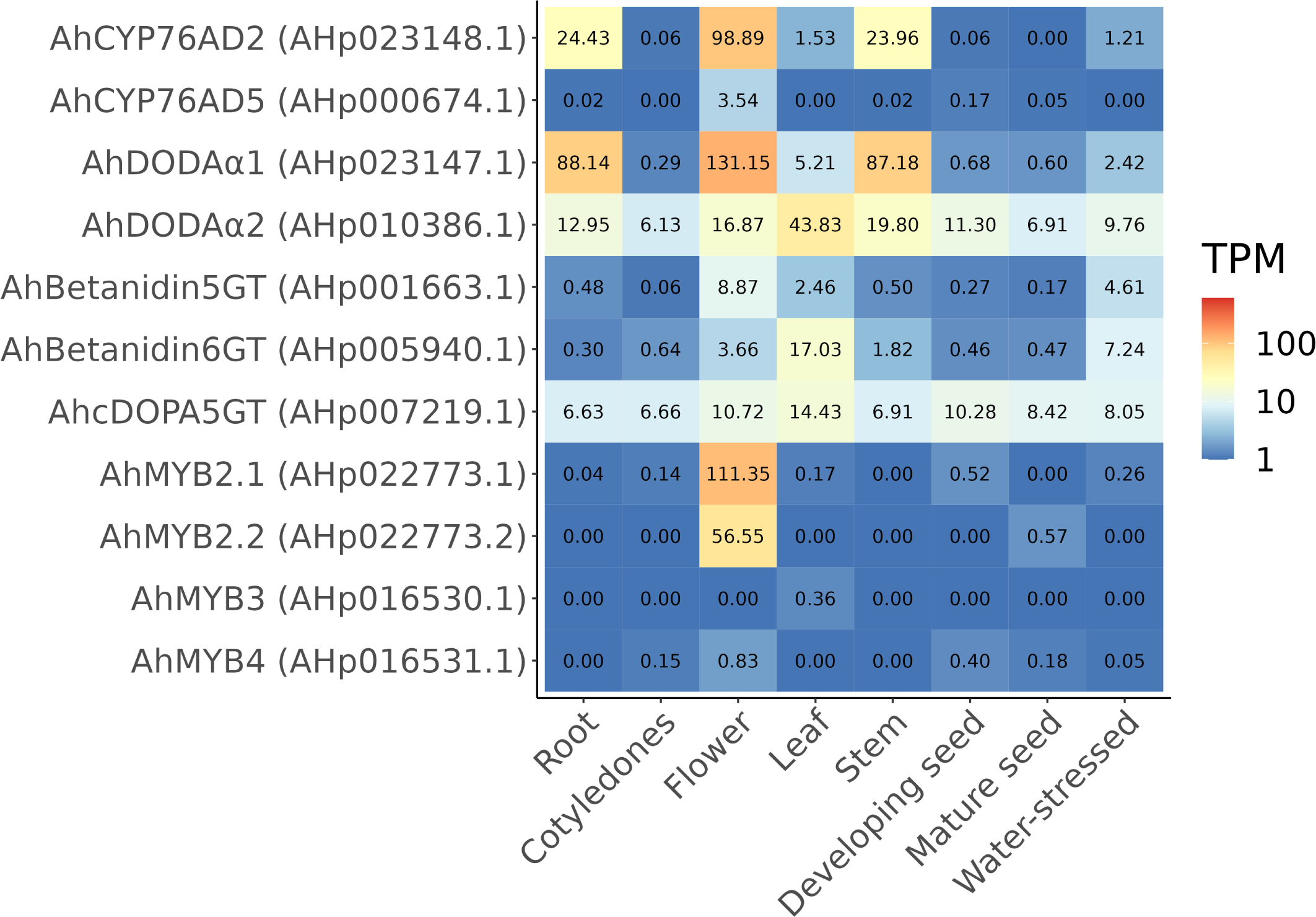
Relative gene expression in TPM of identified betalain pathway gene expression in different tissues. Quantification was done using kallisto from short-read RNA sequencing data.

**Figure S6.**
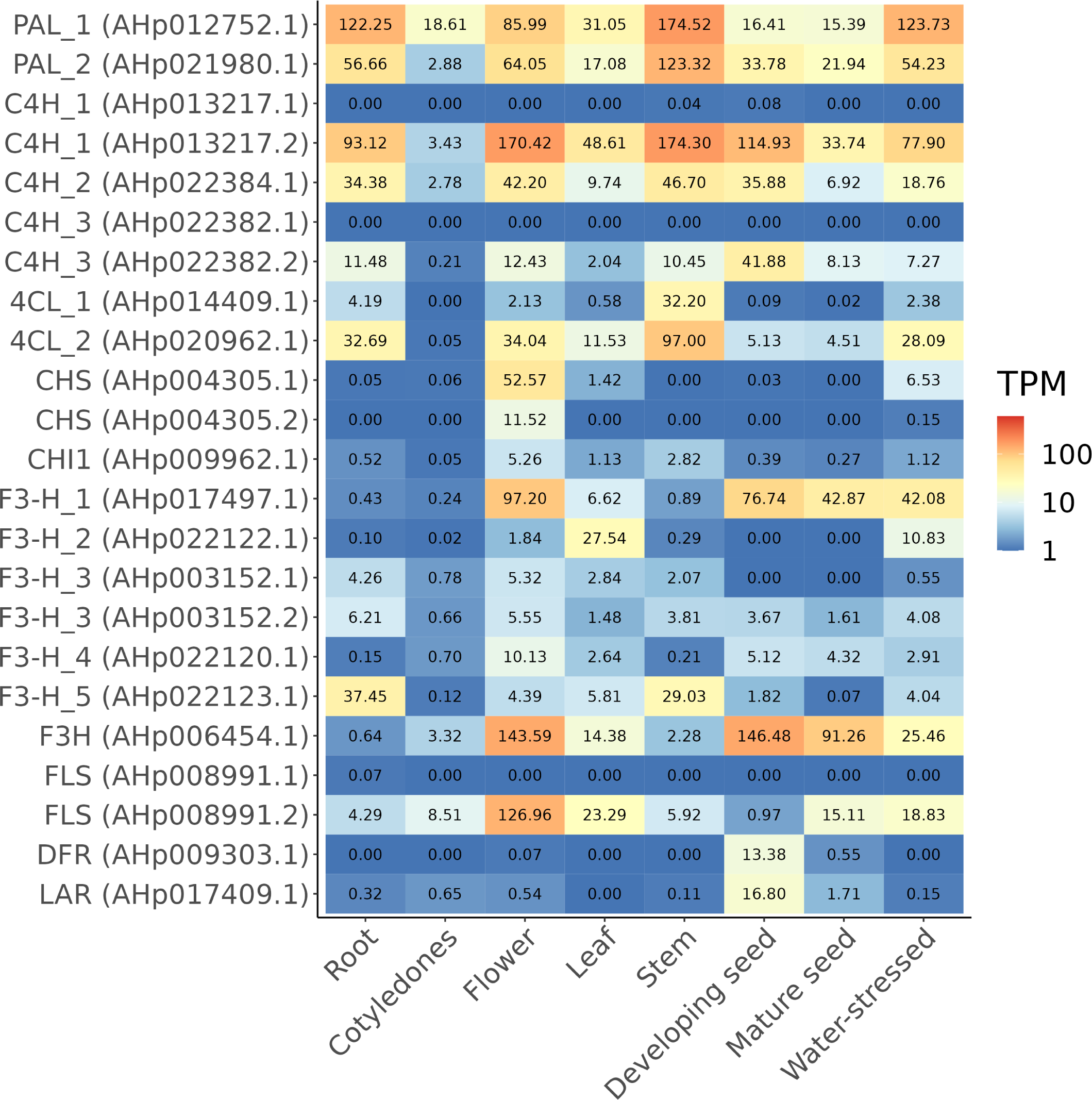
Relative gene expression in TPM of identified flavonoid pathway gene expression in different tissues. Quantification was done using kallisto from short-read RNA sequencing data.

**Figure S7.**
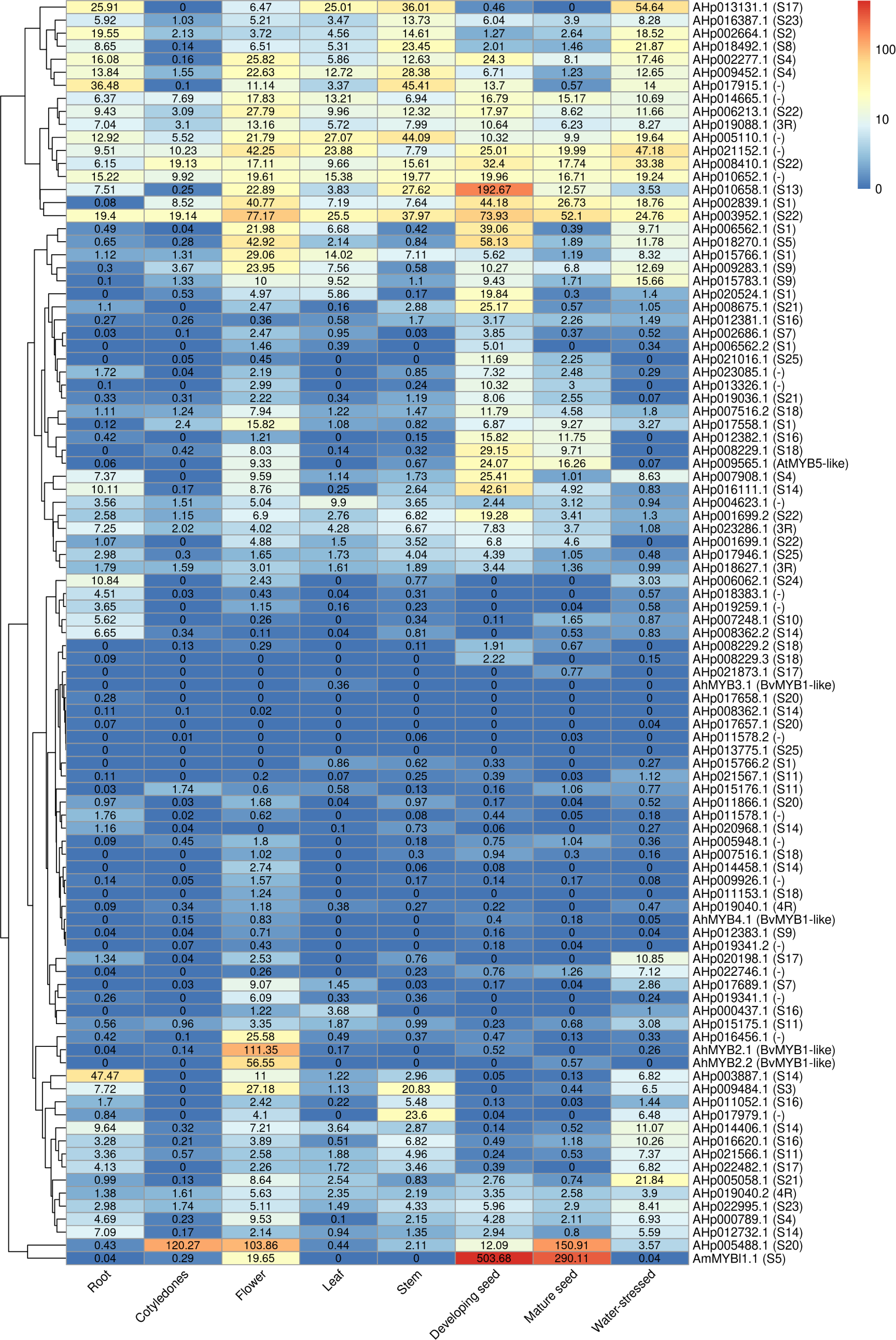
Relative gene expression in TPM of identified MYB transcription factor genes in different tissues. Quantification was done using kallisto from short-read RNA sequencing data.

**Figure S8.**
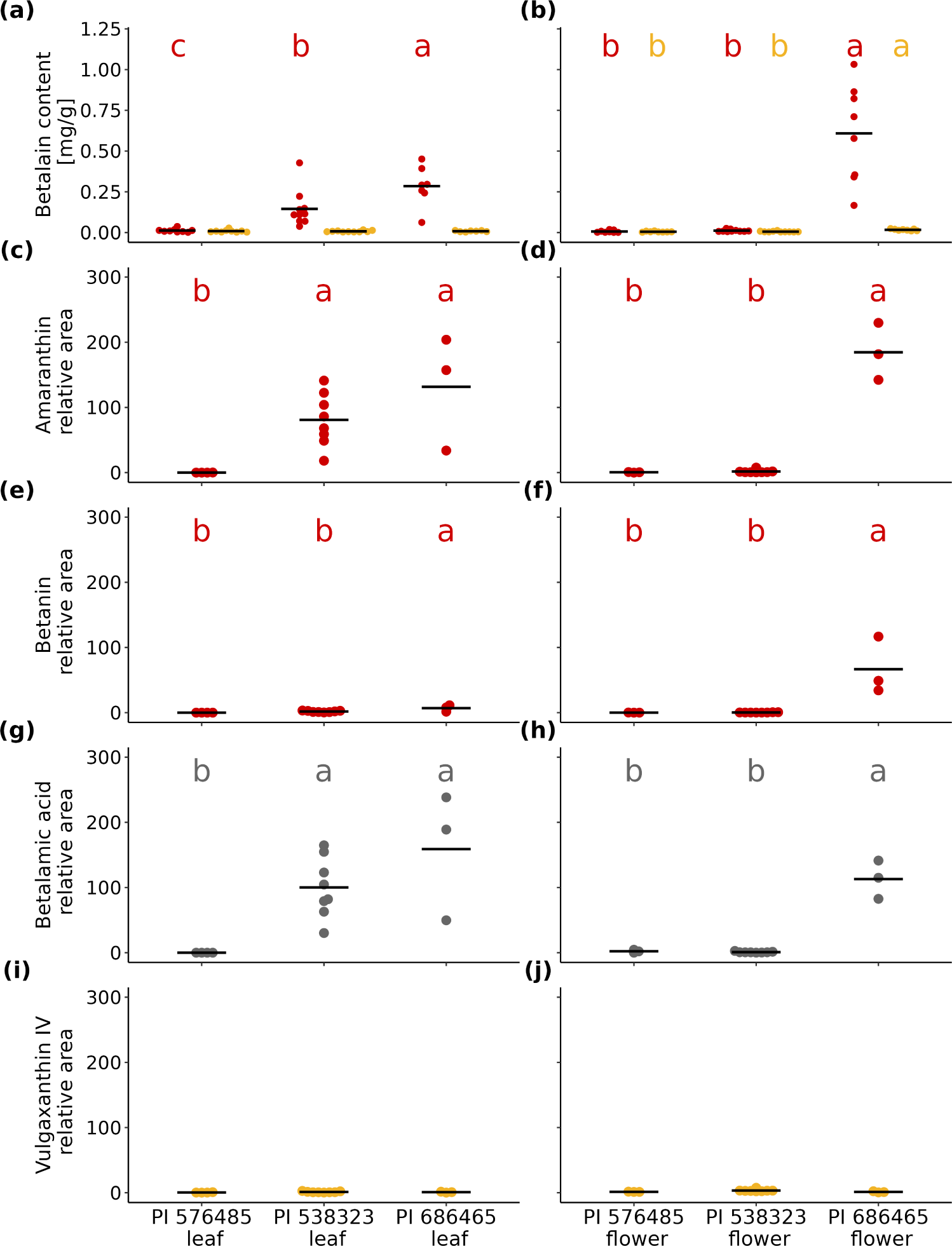
Photometric and LC-MS quantification of betalain pathway metabolites for BSA parental genotypes. (a-b) Photometric quantification of betacyanin (red) and betaxanthin (yellow) content of (a) leaf and (b) flower tissue (mg betalains per g fresh weight), with black bars denoting mean values. (c-f) LC-MS quantification of the betacyanins amaranthin (c, d) and betanin (e, f) in leaf and flower, respectively. (g-h) LC-MS quantification of betalamic acid as common chromophore of betacyanins and betaxanthins in (g) leaf and (h) flower. (i-j) LC-MS quantification of the betaxanthin vulgaxanthin IV in (i) leaf and (j) flower. Statistical analysis was conducted using ANOVA and significant differences in betaxanthin or betacyanin content between accessions within tissues were denoted using compact letter display. There were no significant differences in total betaxanthin content between accessions in leaf and no significant differences in vulgaxanthin IV content between accessions in leaf or flower.

**Figure S9.**
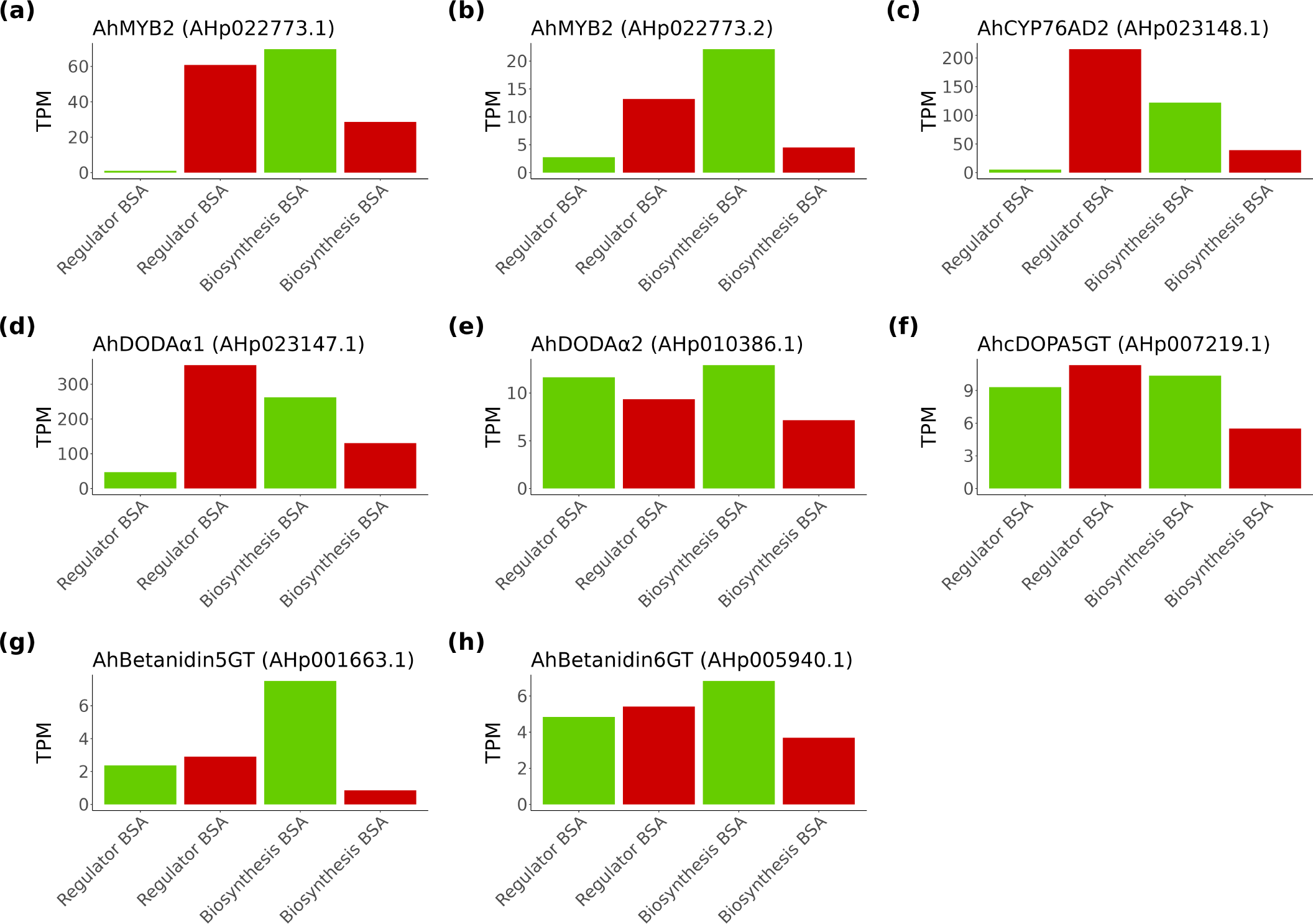
Gene expression of betalain pathway genes in flower tissue in green and red bulks of regulator and biosynthesis BSAs. Relative gene expression in TPM is displayed for betalain pathway genes and candidate regulators. *AhCYP76AD5*, *AhMYB3* and *AhMYB4* had expression < 0.5 TPM in flower tissue in all samples and were excluded.

**Figure S10.**
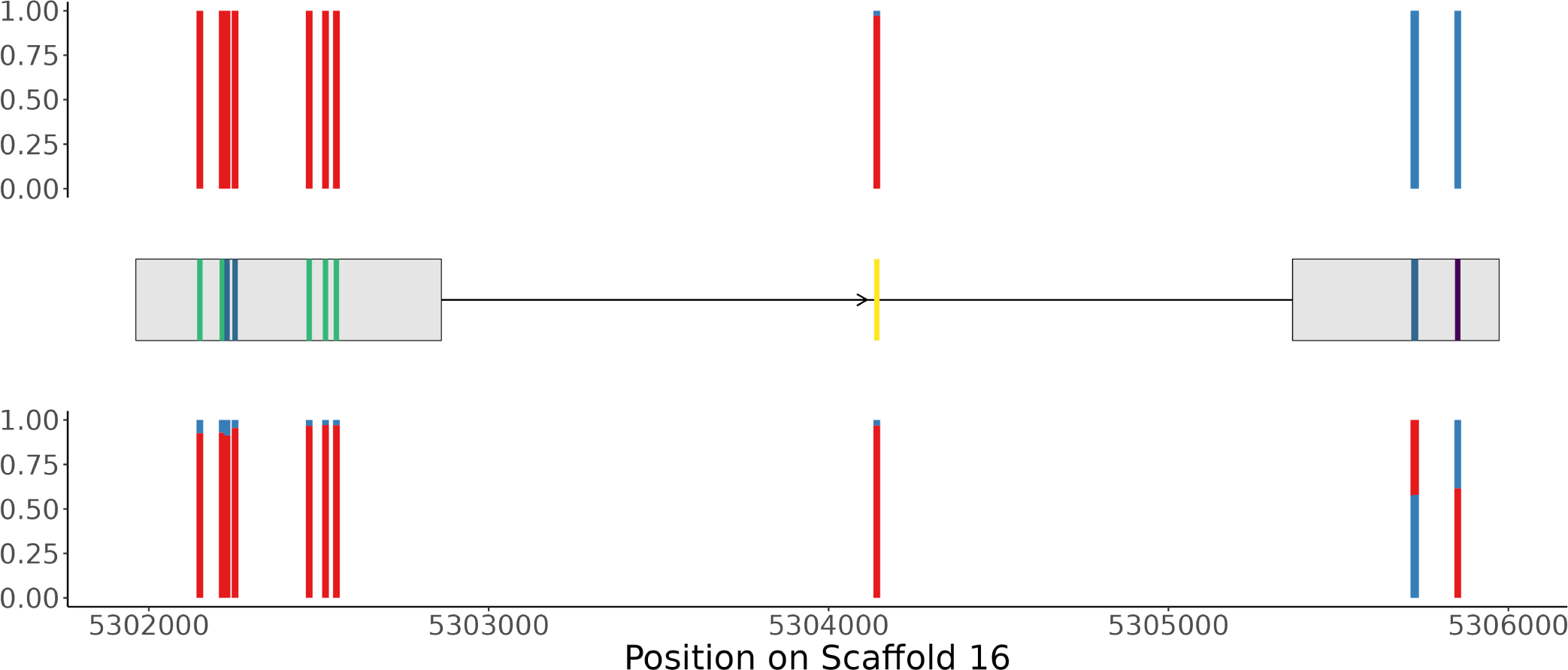
Segregating variants called from whole-genome sequencing data in the biosynthesis BSA and their relative allele depth for the *AhCYP76AD2* gene. Top row shows the relative depth of reference (blue) and alternative allele (red) for the red bulk, the bottom row the relative depth for the green bulk. Middle part shows the exon structure of the gene and all variants with their predicted effect compared to the reference genome (yellow: intron variant, green: synonymous variant, dark blue: missense variant, purple: stop-gain variant).

**Figure S11.**
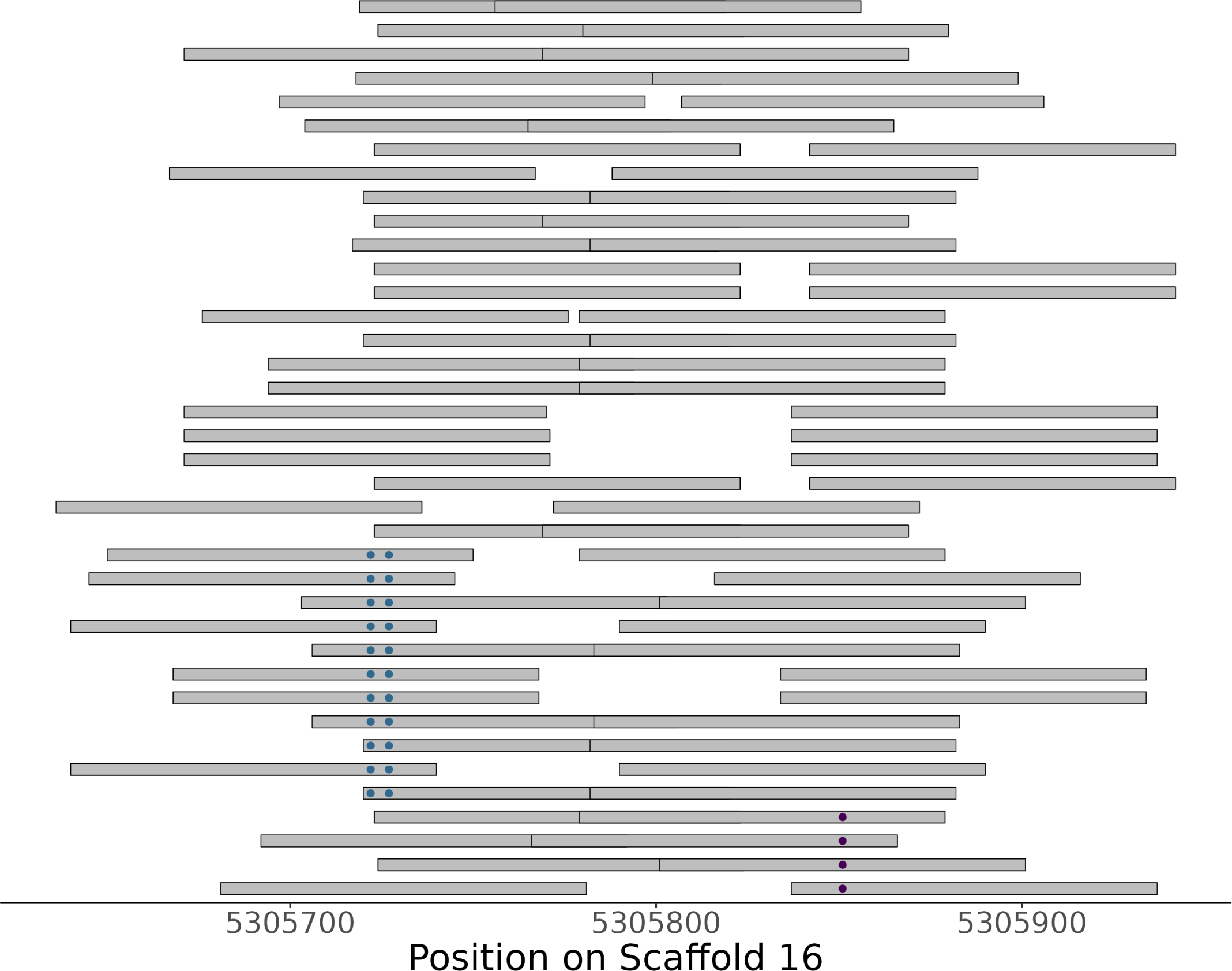
Missense and stop alleles in *AhCYP76AD2* from RNA-seq reads of red flower tissue. RNA sequencing read pairs from flower tissue of the red bulk of the biosynthesis BSA covering at least one of the non-synonymous allele T1258A and A1263T positions and the stop allele C1387T position on the *AhCYP76AD2* gene. Variant positions were colored based on the assigned impact of the allele on a read on the translated protein.

**Figure S12.**
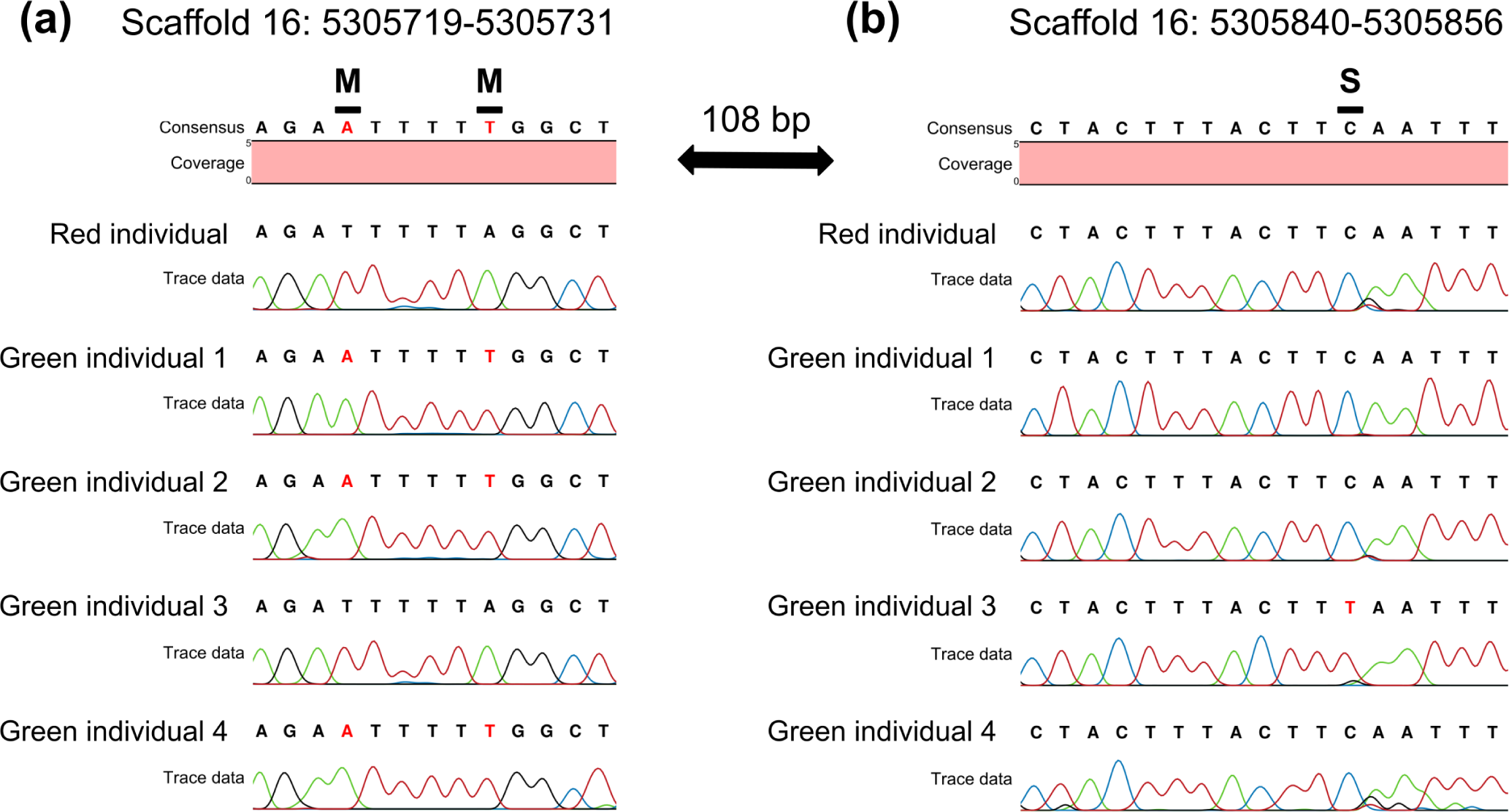
Sanger sequencing of *AhCYP76AD2* from biosynthesis BSA individuals. Sanger sequencing of *AhCYP76AD2* (primer SV4) from individuals from biosynthesis BSA were mapped to the *AhCYP76AD2* genomic sequence. (a-b) One individual was chosen from the red bulk (top), the other four individuals were from the green bulk. Positions differing from the reference genome were marked in red. (a) Positions of the two missense variants T1258A and A1263T were marked with M, (b) the stop variant C1387T was marked with S.

## Notes

### Competing Interest Statement

The authors have declared no competing interest.

### Summary of Updates

- Adjusted the title to: "Isoform-resolved genome annotation enables mapping of tissue-specific betalain regulation in amaranth" - Added description of additional experiments to quantify betalain content in leaf and flower for parental accessions used in mapping and to functionally validate the flower-specific betalain regulator AhMYB2 to methods, results, discussion and supplementary methods S6 - Modified text to improve readability - Modified Figure 1 to include a simplified representations of (a) the general phenylpropanoid and flavonoid pathway and (b) the betalain pathway, modified the circos plot to include the positions of identified pathway genes - Modified Figure 2 to include functional assignments for MYB transcription factor subgroups - Added Figure 3 visualising the parental accessions used for mapping as well as photometric and LC-MS betalain quantification from leaf and flower of the parental accessions - Added Figure 6 showing the functional validation of the flower-specific betalain regulator AhMYB2 and nuclear localisation - Added Figure S8 summarising the results of both, photometric quantification of total betalain content and LC-MS quantification of the betacyanins amaranthin and betanin, the central betalain chromophore betalamic acid and the betaxanthin vulgaxanthin IV

